# Netupitant Exhibits Potent Activity on *Mycobacterium tuberculosis* Persisters

**DOI:** 10.1101/2024.10.30.620736

**Authors:** Hassan E. Eldesouky, Richard M. Jones, Natalie Gleason, Shabber Mohammed, Enming Xing, Pui-Kai Li, David R. Sherman

## Abstract

In *Mycobacterium tuberculosis* (Mtb), persisters are genotypically drug-sensitive bacteria that nonetheless survive antibiotic treatment. Persisters represent a significant challenge to shortening TB treatment and preventing relapse, underscoring the need for new therapeutic strategies. In this study, we screened 2,336 FDA-approved compounds to identify agents that enhance the sterilizing activity of standard anti-TB drugs and prevent the regrowth of persisters. Netupitant (NTP), an FDA-approved antiemetic, emerged as a promising candidate with bacteriostatic activity on its own. However, in combination with isoniazid (INH) and rifampicin (RIF), NTP eliminated viable Mtb cells within 7 days, achieving a >6-log reduction in colony-forming units (CFUs) compared to the 2.5-log reduction observed with INH-RIF alone. NTP also demonstrated broad-spectrum efficacy, enhancing the activity of multiple TB drugs, including ethambutol, moxifloxacin, amikacin, and bedaquiline. Notably, NTP retained its potency under hypoxic and caseum-mimicking conditions, both of which are known to enrich for non-replicating, drug-tolerant cells. Interestingly, under hypoxic conditions, NTP demonstrated strong tuberculocidal activity, achieving an approximate 4-log CFU reduction, whereas high-dose INH-RIF was ineffective. Transcriptomic analysis revealed that NTP primarily disrupts cellular bioenergetics, with significant downregulation observed in activities associated with the electron transport chain, oxidative phosphorylation, NADH-ubiquinone oxidoreductase, succinate dehydrogenase, and ATP synthesis. While further studies are required to decipher the mechanism of action and resistance profile of NTP, and to assess its *in vivo* efficacy, these findings underscore its potential as a promising adjunct to existing TB therapies.

## Introduction

Tuberculosis (TB), caused by the bacillus *Mycobacterium tuberculosis* (Mtb), remains one of the deadliest infectious diseases globally. Despite significant advancements in healthcare, TB continues to be responsible for approximately 1.3 million deaths annually, making it the leading cause of mortality from a single infectious agent. Latent TB infections are very widespread, and there were around 10 million new cases of active TB in 2022 alone (World Health Organization). The disease poses a significant threat to global health, particularly in areas with high rates of HIV co-infection and poor access to healthcare. Furthermore, the COVID-19 pandemic disrupted public health services worldwide, leading to the first rise in TB-related deaths in over two decades (World Health Organization).

Current TB treatment regimens span four to six months and require the use of multiple drugs. The treatment has several limitations, including high toxicity, and long duration. An additional concern is the failure of antibiotics to fully eradicate infection in all cases, even after extended treatment times. In drug-sensitive TB, treatment is generally six months, while in drug-resistant cases, it can extend up to two years ^1^. Around 5-10% of patients fail to respond fully to the initial therapy or experience relapse ^2-4^. These poor outcomes have been associated with a drug-tolerant subpopulation of Mtb cells, commonly referred to as “persisters” ^5^. These cells are poorly eradicated by conventional TB therapies, leading to treatment failure and infection relapse. Furthermore, recent studies have indicated that Mtb persisters contribute to the emergence of drug resistance ^6-9^, thus underscoring the need for new treatments targeting this subpopulation.

In light of these challenges, there has been a growing interest in drug repurposing as a strategy to combat TB. Drug repurposing involves identifying new therapeutic uses for existing drugs, thereby reducing the time and cost associated with developing novel treatments from scratch ^10, 11^. Repurposed drugs often have established safety and efficacy profiles, which can streamline clinical development and potentially lead to faster approval for TB treatment. Moreover, repurposed drugs may target novel mechanisms of action that could overcome resistance to traditional TB medications, thereby expanding the therapeutic options available for patients, particularly those with drug-resistant strains.

In this study, we developed a screening assay to identify compounds capable of targeting Mtb persisters that survive high concentrations of first-line TB drugs, isoniazid (INH) and rifampicin (RIF). Using this assay, we identified the FDA-approved drug netupitant (NTP), which demonstrated promising activity against Mtb persisters. We then explored the activity of NTP in a variety of disease relevant conditions and opened studies to understand NTP action on Mtb. The identification of NTP as a potential anti-TB agent offers a potential new avenue for improving TB treatment outcomes and reducing the duration of therapy.

## Results

### Identification of netupitant (NTP) as an anti-persister agent

To identify small molecules that enhance the sterilizing activity of current anti-TB drugs and inhibit the regrowth of TB persisters, we screened a drug library comprising 2,336 FDA-approved compounds at a fixed concentration (40 μM) against Mtb (H37Rv) cells. The screening was performed with or without high doses of the first-line agents isoniazid (INH) and rifampicin (RIF), simultaneously administered at concentrations of 3.12 µg/ml and 0.4 µg/ml, respectively. These doses correspond to 100 times the minimum inhibitory concentration (MIC) as determined in a standard MIC assay (Supplementary Table 1). After 7 days of incubation at 37°C, cells were washed, resuspended in drug-free 7H9/OADC medium, and cell viability was assessed using the BacTiter glo assay (Fig. 1a). We were able to identify a few compounds that exhibited the ability to enhance the antibacterial activity of INH-RIF. Among those compounds, netupitant (NTP), an antiemetic, emerged as a promising candidate. When combined with INH-RIF, NTP reduced ATP levels by more than 90% compared to INH-RIF treatment alone (Fig. 1b), and even after removal of the drug pressure ATP levels remained extremely low, consistent with cell death and sterilization of the Mtb culture (Fig. 1c). In contrast, we observed a substantial rebound in ATP levels in the INH-RIF group only following the removal of the drug pressure, suggesting cell growth recovery (Fig. 1c). To further validate NTP’s efficacy, time-kill studies and spot assays were performed. Our results showed that NTP exhibited moderate antitubercular activity on its own, displaying a bacteriostatic effect at 16 or 32 μg/ml. However, when combined with high dose INH-RIF, NTP exhibited potent bactericidal activity, resulting in a greater than 6-log reduction in CFU within 7 days, compared to only a 2.5-log reduction with INH-RIF alone (Fig. 1d). These findings were further corroborated by spot assays showing survival CFU in response to the respective treatments (Fig. 1e).

**Table 1.**
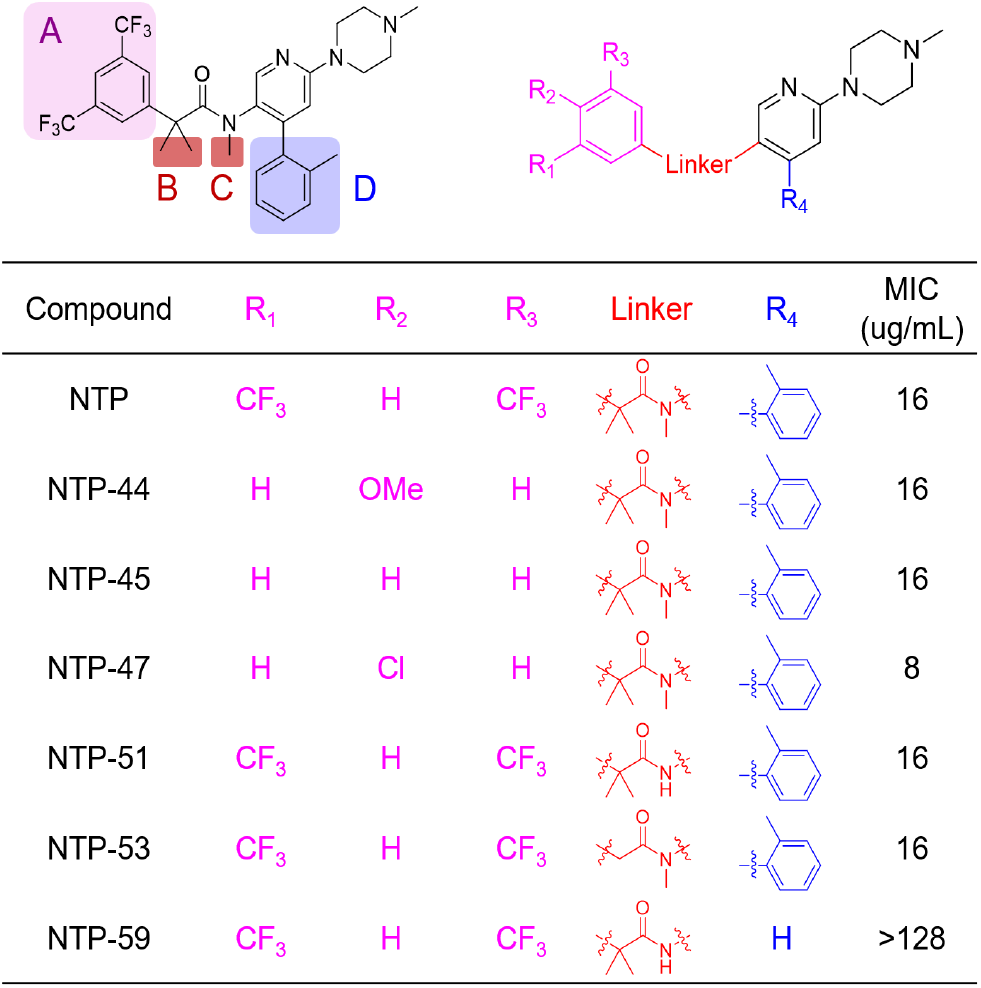
The preliminary structure-activity relationship table for NTP analogs, with Mtb MICs.

**Figure 1:**
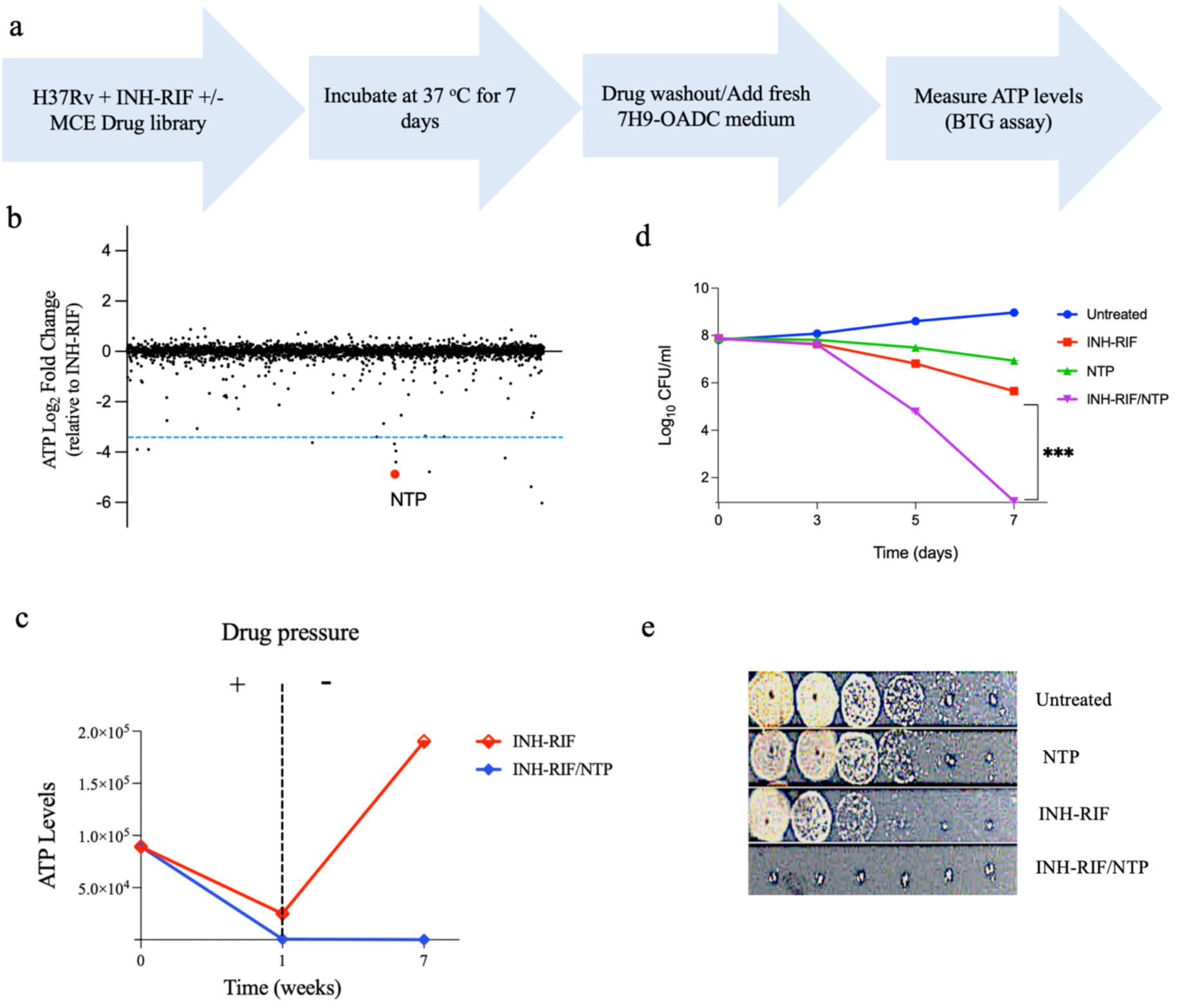
Identification of netupitant (NTP) as an anti-persister agent in Mtb. (a) Schematic diagram illustrating the screening of the drug library against the *M. tuberculosis* H37Rv strain in the presence or absence of high dose INH-RIF (3.12/0.4 μg/ml). (b) Library screening results measuring ATP levels in H37Rv cells after drug removal, followed by incubation in drug-free medium for 6 weeks. A dashed blue line was added to indicate a 90% depletion in ATP levels, corresponding to a -3.32 log_2_ fold change relative to the INH-RIF control. (c) Seven-week time course showing temporal measurements of ATP levels in H37Rv cells treated with high dose INH-RIF ± NTP (2x MIC), followed by removal of drug pressure. (d) Time-kill curve illustrating the effect of a 7-day treatment with NTP (2x MIC) in the presence or absence of high dose INH-RIF on H37Rv cells, with asterisks indicating statistical significance (*p*<0.05). (e) Spot assay demonstrating H37Rv colony growth on 7H10 agar plates following 7 days of treatment with NTP (2x MIC) ± high dose of INH-RIF.

### Interactions with standards anti-TB drugs

To explore potential interactions between NTP and a range of antitubercular drugs, we conducted checkerboard assays to assess the combined effects of NTP with six commonly used anti-TB agents. Checkerboard results indicated that NTP displayed indifferent interactions with both INH and RIF, with fractional inhibitory concentration indices (ΣFICI) values of 1 and 2, respectively (Figure 2a, b). Furthermore, similar indifferent interactions were observed between NTP and ethambutol (EMB), moxifloxacin (MOX), and amikacin (AMK), each yielding an ΣFIC value of 2. Of particular interest, however, we identified a potent synergistic interaction between NTP and bedaquiline (BDQ), with a ΣFIC value of 0.375 (Figure 2a, b).

**Figure 2:**
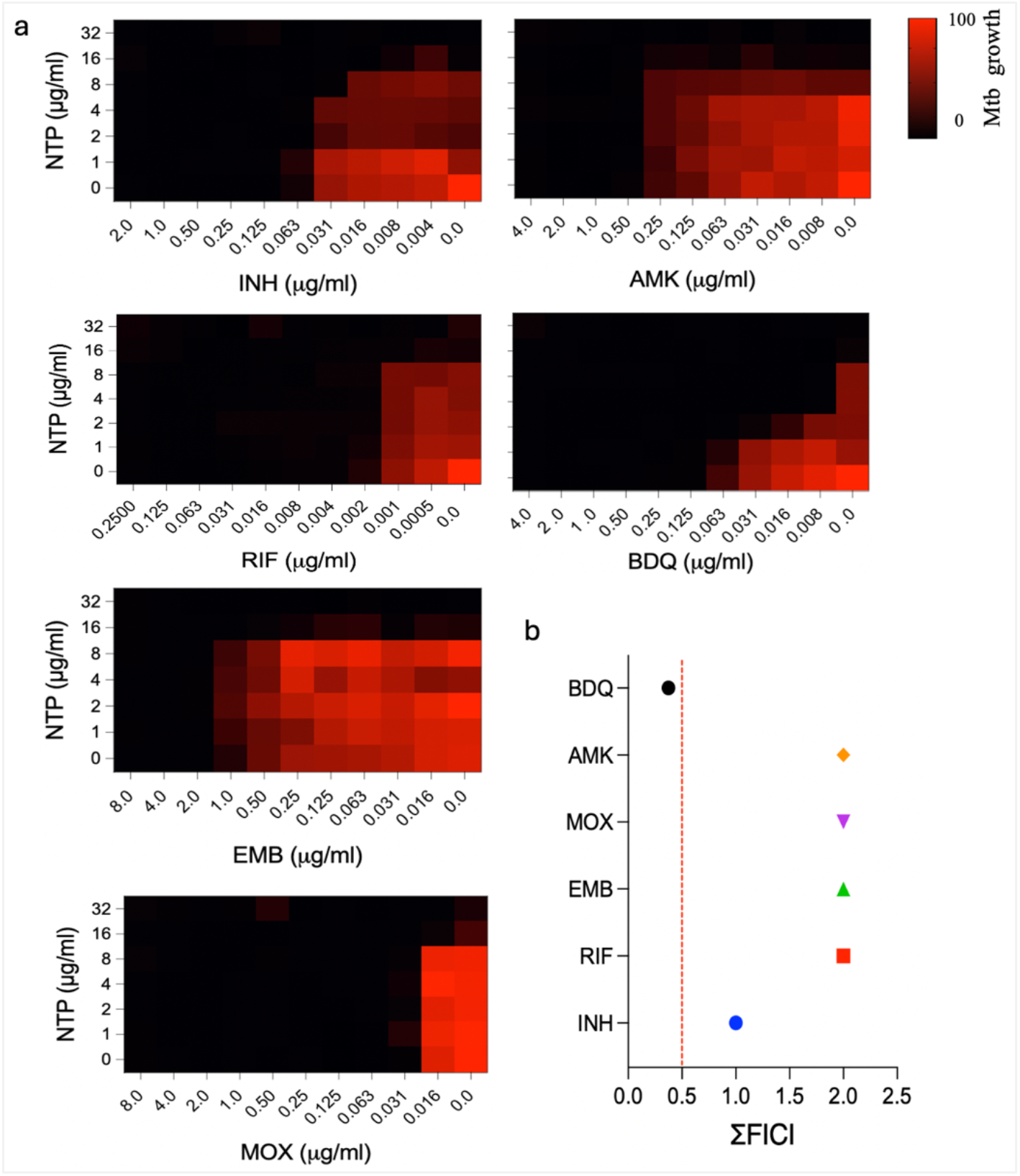
Interactions between NTP and anti-TB drugs (checkerboard method). (a) Checkerboard heatmaps showing the effect of NTP (0–32 μg/ml) on the growth inhibitory activities of isoniazid (INH), rifampicin (RIF), ethambutol (EMB), moxifloxacin (MOX), amikacin (AMK), and bedaquiline (BDQ). Activities measured by OD. (b) Fractional inhibitory concentration indices (ΣFICI) calculated from the checkerboard data, with a dashed red line indicating 0.5 as the threshold for synergy

**Figure 3:**
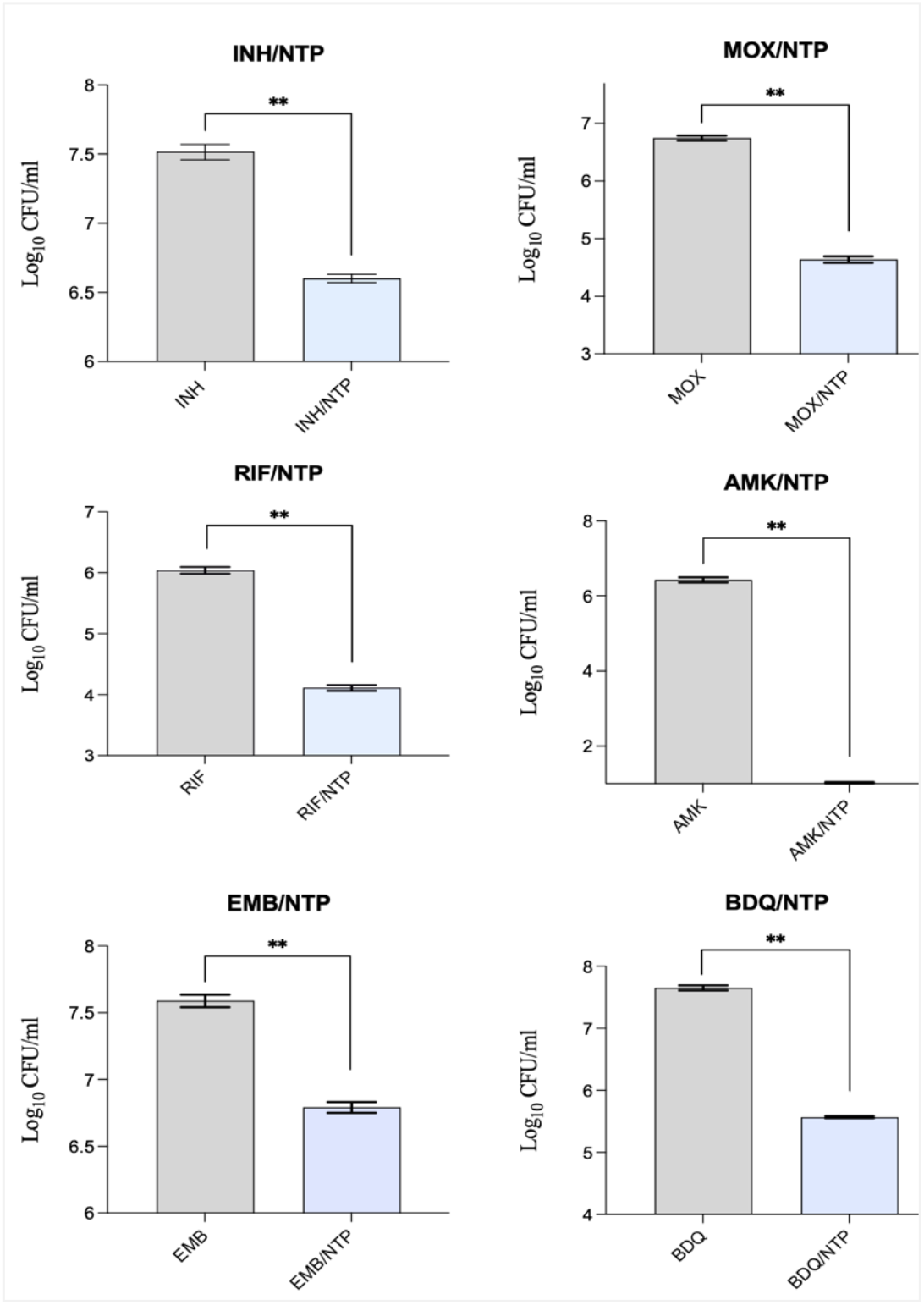
NTP improves the killing activity of anti-TB drugs. H37Rv cells were treated for 14 days with NTP (2x MIC) and 100x MIC of isoniazid (INH), rifampicin (RIF), ethambutol (EMB), moxifloxacin (MOX), amikacin (AMK), and bedaquiline (BDQ). Data are presented as log_10_ CFU/ml, with asterisks indicating statistical significance (*p*<0.05).

### NTP improves the killing activity of a broad spectrum of anti-TB drugs

Our screening results demonstrate that NTP enhances the bactericidal activity of the INH-RIF combination. To further investigate whether NTP’s activity exhibits specificity towards certain antitubercular drugs, we conducted a 14 day time-kill study to assess its interaction with a range of current anti-TB agents. We examined the effect of NTP (32 ug/ml) on INH, RIF, EMB, MOX, AMK, and BDQ, tested at concentrations equivalent to 100x MIC in standard MIC assays (Supplementary Table 1). Our time kill kinetic data demonstrated that NTP was able to improve the killing efficacy of all tested drug. When combined with either RIF, MOX, or BDQ, NTP resulted in an additional reduction in viable CFU by 2 ±0.08 logs compared to monotherapy with each drug. The combination of NTP and AMK demonstrated an even greater effect, achieving a 5.7 ± 0.7 log reduction in CFU counts relative to AMK alone. Additionally, we found that NTP enhanced the killing activity of INH and EMB, leading to CFU log reductions of 0.9 ± 0.03 and 0.79 ± 0.04, respectively, compared to treatment with INH or EMB alone These data indicate that NTP exhibits broad-spectrum activity when combined with various antitubercular drugs, potentially targeting a universal tolerance mechanism in Mtb.

### NTP activity against non-replicating TB

A significant challenge in TB treatment lies in the capacity of Mtb cells to enter a drug-tolerant state. This state can be induced by various stressors, both in vitro and in vivo, such as drug pressure, starvation, hypoxia, and residence in caseous granulomas^12-14^. To assess whether NTP retains its activity in various conditions conducive to drug tolerant Mtb, we evaluated its effect in the presence or absence of INH-RIF in hypoxic and caseum surrogate models. Both conditions are reportedly associated with drug tolerance and Mtb cells are not able to grow in these environments.

In caseum-mimicking conditions, NTP alone did not exhibit a substantial killing effect. However, high dose INH-RIF treatment reduced CFUs by 3 logs by day 21 (Figure 4a). The addition of NTP to INH-RIF further enhanced its bactericidal activity, resulting in 6.3 ±0.07 log reduction in viable CFU, relative to the untreated control (Figure 4a).

**Figure 4:**
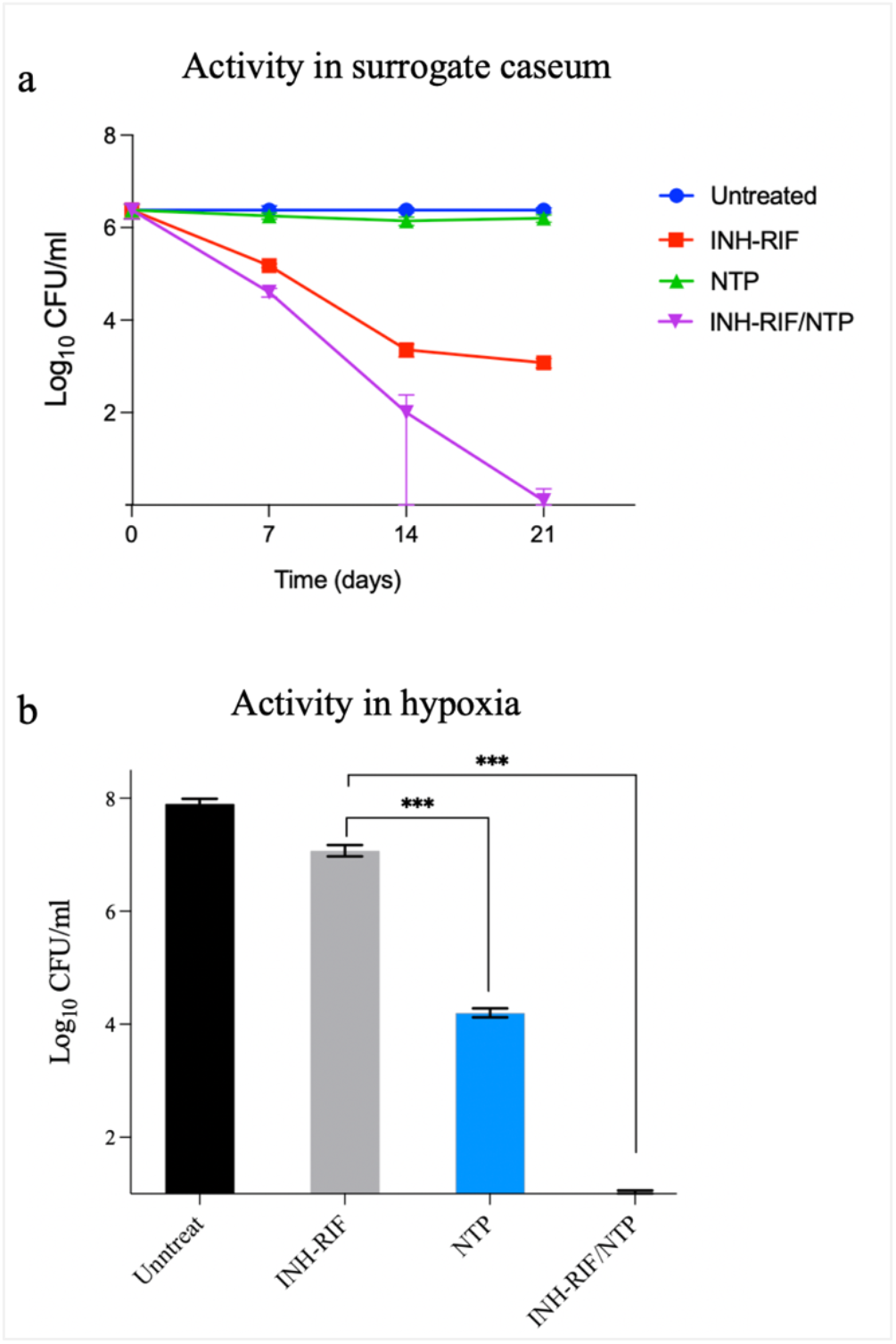
Activity of NTP against non-replicating Mtb cells. (a) CFU analysis of H37Rv cells treated with NTP (32 ug/ml) ± high dose INH-RIF for 21 days in surrogate caseum. Asterisks indicates statistical significance (*p*<0.05). (b) CFU analysis of H37Rv cells treated with NTP (32 ug/ml) ± high dose INH-RIF for 14 days under hypoxic conditions is a GasPak

In hypoxic conditions, treatment with high dose INH-RIF resulted in only a modest reduction of viable CFUs, with a 0.8 log reduction after 14 days. However, the addition of NTP to the INH-RIF regimen led to a nearly 7 log reduction in CFUs (Figure 4b), with no viable cells detectable above the limit of detection (10 CFU/ml). Notably, in contrast to its bacteriostatic activity on log-phase Mtb, NTP alone at 2x MIC demonstrated significant bactericidal activity under hypoxic conditions, achieving a 3.7 ±0.06 log reduction in CFUs by day 14 (Figure 4b). Moreover, extending NTP treatment to 30 days resulted in complete eradication of Mtb cells (data not shown).

Additionally, under these hypoxic conditions, the efficacy of NTP treatment surpassed all tested anti-TB drugs, including AMK, EMB, MOX, and BDQ (Supplementary Figure 1). Collectively, these data indicate that NTP significantly enhances the bactericidal activity of INH-RIF against non-replicating Mtb cells under conditions that are unfavorable for most standard anti-TB drugs.

### Structure Activity Relationship (SAR) analysis

Our structure-activity relationship (SAR) analysis of NTP revealed key insights into its structural determinants of activity. Through systematic modification of specific molecular regions, we identified both essential and non-essential structural elements. The investigation focused on four key structural regions (A, in pink; B, in red; C, in red; and D, in blue) of the NTP framework. At position A, the di-trifluoromethyl groups demonstrated flexibility in modification. Complete removal of these groups (NTP-45) or replacement with single substituents such as methoxy (NTP-44) or chloro (NTP-47) maintained comparable inhibitory activity to the parent compound. Analysis of regions B and C showed that neither the dimethyl groups nor the N-methyl moiety significantly contributed to the compound’s activity. Derivatives NTP-51 and NTP-53, lacking these respective groups, retained activity profiles similar to NTP. The critical structural feature proved to be the tolyl group at position D. Its removal, as demonstrated in compound NTP-59, resulted in complete loss of activity, establishing this moiety as essential for the molecule’s biological function. Detailed information on the synthesis of NTP and its analogs is provided in the supplementary document (Supplemental Materials 2.docx).

### Transcriptomic Profiling Reveals NTP’s Dynamic Impact on *M. tuberculosis* Metabolic Pathways

To investigate the molecular mechanisms underlying NTP’s observed activity, we conducted RNA-seq analysis on Mtb H37Rv at 1, 6, and 24 hours post-treatment with NTP. We observed that NTP triggered a rapid and extensive transcriptional response. The number of differentially expressed genes (DEGs), defined as those with at least a 2-fold change compared to the untreated control (p < 0.05), ranged from 297 to 622 upregulated genes and 263 to 1021 downregulated genes across the time points. Gene set enrichment analysis revealed that NTP powerfully disrupted cellular bioenergetics. Downregulated pathways included those involved in the electron transport chain (ETC), oxidative phosphorylation, NADH-ubiquinone oxidoreductase, succinate dehydrogenase, and ATP synthesis (Fig. 5a). In contrast, central carbon metabolic processes, such as pyruvate dehydrogenase and 3-methyl-2-oxobutanoate dehydrogenase, were upregulated, possibly as a compensatory response to the impaired ETC function. Aligned with these findings, we observed that NTP alone significantly reduces ATP levels in a dose dependent manner, compared to the untreated control (Supplementary Figure 2). We also observed a significant upregulation in genes related to iron-sulfur cluster assembly, which are crucial for the functionality of several ETC components. Additionally, pathways related to oxidative stress response, protein metabolism, and nucleic acid metabolism were broadly downregulated, highlighting the extensive impact of NTP on multiple essential cellular processes (Fig. 5a).

**Figure 5:**
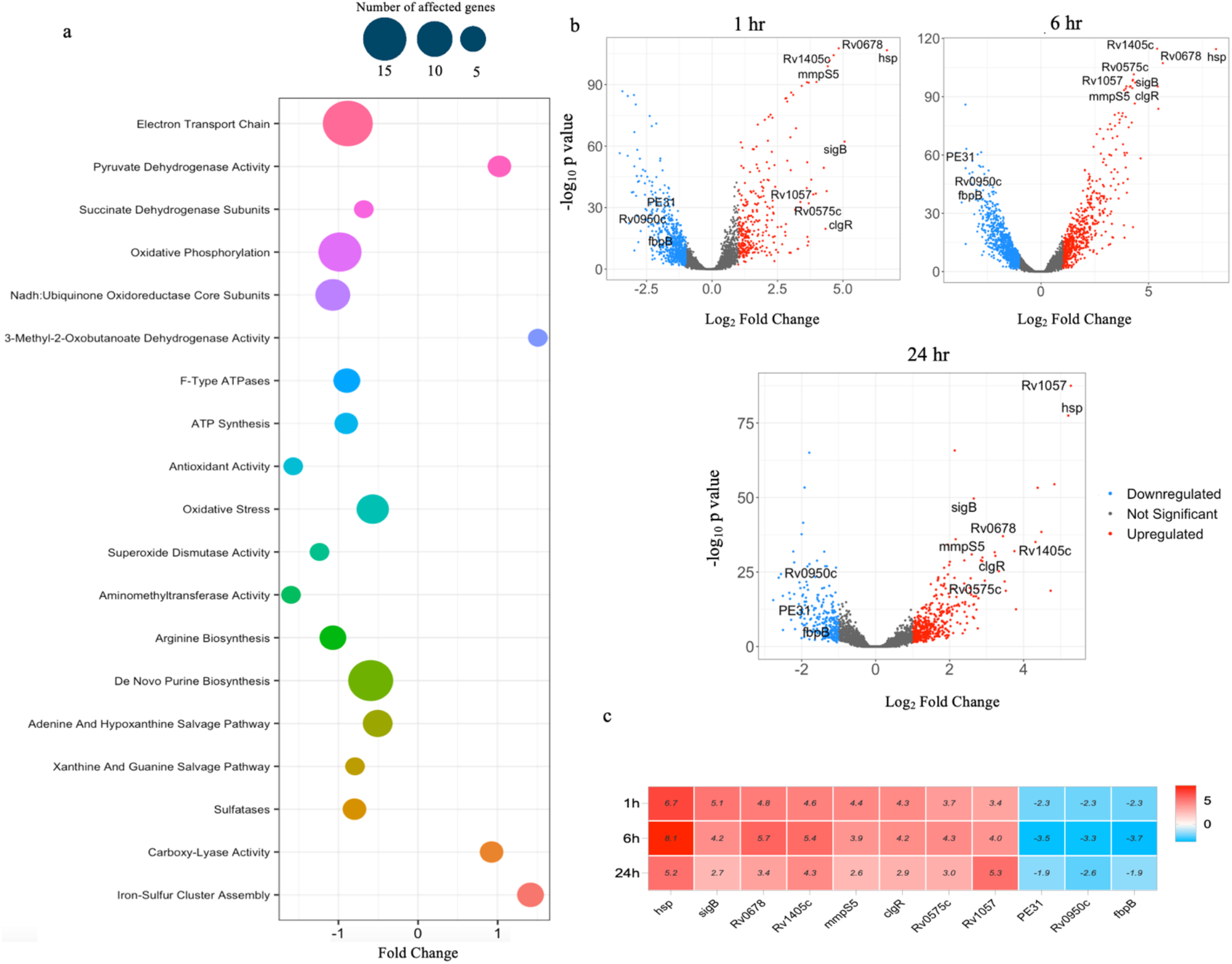
Transcriptomic profiling of NTP’s effect on Mtb. (**a**) Gene Ontology (GO) terms representing biological processes that were significantly enriched (p < 0.05) across all time points in response to NTP treatment, highlighting both over- and underrepresented processes. (**b**) Volcano plots showing the impact of NTP (32 ug/ml) on the H37Rv transcriptome at 1, 6, and 24 hours. The plots depict the significantly differentially expressed genes (p < 0.05) with at least a 2-fold change compared to the untreated control. (**c**) Heatmap displaying the expression patterns of the top 11 genes that showed consistent regulation across all time points.

To further investigate NTP’s effect on Mtb, we explored individual transcriptional responses. We identified the top 100 most-regulated genes at each time point and found 11 genes that consistently exhibited significant changes in expression throughout the study. These genes are highlighted in the volcano plots and heat maps of Fig. 5b and c. Rv0251 (*hsp*) was the most upregulated gene relative to the control. Previous studies have shown that *hsp* is highly expressed in various non-replicative persistence models, such as the hypoxic Wayne model, nutrient starvation, drug persister models, and in the phagolysosomes of activated macrophages^15^. Additionally, Rv0678 and mmpS5 were among the top upregulated genes. Rv0678 is a known repressor of the mmpL5-mmpS5 transporter system, which plays key roles in efflux activity and iron homeostasis ^16-19^. The co-upregulation of Rv0678 and a gene that it represses suggests that NTP may bypass Rv0678 to exert direct action on the mmpL5-mmpS5 system. To explore this hypothesis, we performed an ethidium bromide efflux assay and evaluated the effect of iron supplementation on NTP’s activity. Our results demonstrated that NTP did not interfere with ethidium bromide efflux, nor was its activity affected by the addition of ferrous sulfate (Supplementary Figure 3), suggesting that NTP’s mechanism of action is independent of mmpL5-mmpS5 activity, though this efflux system likely plays a role in NTP resistance.

## Discussion

TB persisters pose a significant challenge in TB therapy due to their association with infection relapse, prolonged treatment, and the emergence of drug-resistant isolates. These persister cells exhibit tolerance to anti-TB drugs primarily because of their slowed growth and metabolic adaptation, which protects them from the bactericidal effects of antibiotics that target actively replicating bacteria ^5, 20-22^. As such, there is an urgent need to discover new agents that can target persister cells, particularly under conditions where traditional anti-TB drugs are less effective. In this study, we identified netupitant (NTP), an FDA-approved antiemetic, as a potent anti-persister agent that enhances the sterilizing activity of the two first line anti-TB drugs, isoniazid (INH) and rifampicin (RIF), preventing the regrowth of Mtb persister cells. Time-kill studies further demonstrated that NTP exhibited broad spectrum killing-enhancing activities when combined with other anti-TB drugs such as ethambutol (EMB), moxifloxacin (MOX), bedaquiline (BDQ), and amikacin (AMK). This suggests that NTP works by targeting a universal mechanism of tolerance in Mtb, an essential feature for potential therapeutic agents aimed at tackling persister cells.

We also observed potent activity of NTP in hypoxic and caseum-like conditions, which are known to foster drug-tolerant Mtb populations. Hypoxic and caseous environments are both observed in TB patients with granuloma and cavitary lesions, and their presence is associated with treatment failure and disease relapse ^23^. Previous studies have demonstrated that Mtb persisters, which survive under these hostile environmental conditions, contribute to relapse and treatment failure ^2, 24^. Our data indicate that NTP exhibits potent bactericidal activity under hypoxic conditions, whereas all tested anti-TB drugs, even at high doses, showed no activity. This finding could be clinically significant, suggesting that NTP targets an essential pathway under non-replicative conditions. By targeting these persister populations, NTP could play a role in shortening TB therapy and reducing the likelihood of relapse, addressing one of the most significant obstacles in TB treatment.

The observed synergy between NTP and BDQ in our checkerboard assays could provide useful insights into how NTP works. BDQ targets ATP synthase and disrupts energy production in Mtb. NTP’s enhancement of BDQ’s activity suggests that it may potentiate the disruption of Mtb bioenergetics, as indicated by its ability to reduce ATP levels and supported by our transcriptomic data. We observed downregulation of genes involved in the electron transport chain activity, oxidative phosphorylation and ATP synthesis, which may contribute to the bactericidal effects of NTP under hypoxic conditions. Inhibition of bioenergetic pathways could also contribute to NTP’s efficacy even in environments where standard anti-TB drugs are less effective. Notably NTP, treatment induced the co-upregulation of the repressor Rv0678 and its target genes *mmpL5* and *mmpS5*, which are part of the resistance-nodulation-division (RND)-type transporter family^25^. Our in-house data (not shown) suggest that this response also occurs under conditions where cellular bioenergetics are compromised, such as during hypoxia or upon exposure to proton motive force (PMF) inhibitors. Given that the RND efflux system depends on PMF as an energy source^25^, it is plausible that NTP-mediated disruption of the cell bioenergetics, impairs the negative regulatory feedback controlling these transporters.

In humans, NTP exerts its antiemetic effect by selectively blocking the G-protein-coupled receptor (GPCR) NK1, which is involved in vagal signaling linked to nausea and vomiting ^26^. Given that GPCRs are exclusive to eukaryotes, NTP’s mechanism against Mtb likely involves novel bacterial targets that remain to be elucidated. Although the FDA Center for Drug Evaluation and Research has reported that NTP reaches low serum levels in animal models, it also highlighted a marked tendency of NTP to accumulate in pulmonary tissues ^27^, suggesting favorable lung-targeting properties. This pharmacokinetic profile, along with its established safety profile and oral bioavailability, supports NTP potential for repurposing as an anti-TB agent with the goal of reducing treatment duration and improving relapse-free cure rates. Additionally, our preliminary structure-activity relationship (SAR) analysis provides valuable insights for future structural optimization, including the possible development of more potent derivatives.

In conclusion, our findings suggest that NTP holds considerable promise as an adjunct to existing anti-TB therapies, particularly in targeting persister cells and enhancing the activity of TB drugs. Future studies should focus on elucidating NTP’s precise mechanism of action and resistance profile, as well as optimizing NTP-based regimens, including combination therapies that minimize the risk of resistance while maximizing persister eradication. Additionally, testing NTP in animal models of persistence and drug tolerance will be critical for assessing its potential in clinical applications.

## Material and Methods

### Antimicrobials, reagents, and culture conditions

MCE Drug library, netupitant and bedaquiline were purchased from Med Chem Express (NJ, USA), while rifampicin, isoniazid, moxifloxacin, and amikacin were purchased from Sigma-Aldrich (MO, USA). Stock solutions for all compounds except amikacin and isoniazid were prepared in dimethyl sulfoxide (DMSO) and stored at 4°C. Amikacin and isoniazid solution was prepared in water. Actively replicating cultures of *Mycobacterium tuberculosis* H37Rv were grown in Middlebrook 7H9 media (10% OADC [oleic acid, bovine albumin, dextrose, catalase], 0.2% glycerol, 0.05% Tween 80) to an optical density at 600 nm (OD_600_) of 0.5. Cultures were incubated at 37°C on a rotary shaker.

### Minimum inhibitory concentration (MIC) determinations

MIC determinations were performed according to the CLSI protocol M24Ed3 with minor modifications ^28, 29^. Test agents were 2-fold serially diluted in Middlebrook 7H9 OADC/tween/glycerol medium in a volume of 100 μl in 96-well culture plates. Then, aliquots of100 μl of 10^6^ CFU/ml H37Rv strain were added to each well of the drug dilution plates, yielding a final volume of 200 μl. Plates were then incubated at 37°C for 7 days before growth was detected visually and spectrophotometrically at OD_600_. MIC was defined as the minimum concentration that inhibited the growth of Mtb culture by at least 90% relative to the untreated control.

### Drug Screen

*M. tuberculosis* H37Rv strain was cultured in 7H9/Tween/OADC medium to an optical density at 600 nm (OD_600_) of 0.5. Assay plates were prepared by dispensing 75 μL of 7H9 OADC/Tween/glycerol medium into each well of sterile, clear bottom 96-well V-bottom plates (Greiner), supplemented with 40 μM of compounds from the drug library (MCE). All compounds were screened in combination with a final concentration of INH-RIF (3.12/0.4 μg/ml), while control wells contained only INH-RIF. Equal volumes of H37Rv cultures were inoculated into the plates, which were then incubated at 37°C for 7 days. After incubation, plates were centrifuged, supernatants discarded, and wells washed with PBS before adding fresh drug-free 7H9/tween/OADC media. Cell viability was assessed by measuring total ATP levels using the Bactiter Glo assay ^30^.

### Checkerboard assays

Drug-drug interactions were assessed using a two-dimensional (2D) checkerboard titration in 96-well plates, following previously descried protocols ^31-33^. Briefly, anti-TB drugs were serially diluted in 7H9-rich medium and dispensed along the x-axis (50 µL per well), while netupitant was serially diluted along the y-axis (50 µL per well). Mtb H37Rv cultures (OD600 0.005, ∼2×10^6 CFU) were added to each well (100 µL). Plates were sealed in zip-lock bags and incubated at 37°C for 7 days. Cell growth was determined visually and spectrophotometrically at OD_600._ The minimum concentration required to inhibit 90% of growth (MIC90) was determined. Synergy was assessed by calculating the fractional inhibitory concentration (FIC) for each drug. Synergy was defined as ΣFIC ≤ 0.5, antagonism as ΣFIC ≥ 4.0, and no interaction with ΣFIC between 0.5 and 4.0.

### Time-kill assays

Time kill assays were conducted following previously described protocols ^34, 35^. Briefly, Mtb (H37Rv) cultures were grown to mid-log phase (OD_600_ 0.5) in 7H9-rich media. Test agents or vehicle controls were added at the respective concentrations. Samples were collected at designated time points, serially diluted, and plated on 7H10 agar. Colonies were counted after 3 weeks of incubation.

### Activity in surrogate caseum

Surrogate caseum was prepared as described previously ^36^. Briefly, THP-1 cells were cultured in RPMI 1640 with 10% fetal bovine serum and 2 mM l-glutamine, seeded in 150-mm dishes, and differentiated with 100 nM PMA. Lipid accumulation was induced with 0.1 mM stearic acid, and after 24 hours, foamy macrophages were collected, lysed by freeze-thaw cycles, and incubated at 75°C for 30 minutes. The caseum surrogate was rested at 37°C for 3 days before being infected with ∼1×10^8 CFU of the H37Rv strain. The Mtb cells were allowed to adapt to the caseum environment for 6 weeks at 37°C before treatment with NTP (2×MIC) +/-INH-RIF (100×MIC) and re-incubated for 3 weeks. Afterward, surviving CFU were serially diluted in PBS, plated onto 7H10 agar, and counted after 21 days of incubation.

### Activity in hypoxic conditions

Mtb (H37Rv) cultures were grown to mid-log phase (OD600 0.5) in 7H9-rich media. 100 µL aliquots were added to 96-well plates containing test agents or vehicle controls in 7H9-rich medium. The plates were sealed in anaerobic bags (BD GasPak EZ; BD Diagnostics) and incubated at 37°C with 5% CO2 and 100% humidity for 7 days. Samples were then collected, serially diluted, and plated on 7H10 agar and colonies were counted after 3 weeks of incubation.

### RNA sequencing and analysis

RNA was isolated as previously described ^28, 37-39^. Briefly, cell pellets in Trizol were homogenized using Lysing Matrix B and a FastPrep 120 homogenizer, followed by centrifugation. The supernatant was extracted with chloroform and RNA was precipitated with isopropanol and high salt solution. Purification was done using RNeasy kit with DNase treatment (Qiagen), and RNA yield was quantified using a Nanodrop. Ribosomal RNA was depleted using the RiboZero kit (Illumina), and mRNA libraries were prepared with the NEBNext Ultra RNA Library Prep Kit (New England Biolabs) and barcoded using NEBNext Multiplex Oligos. Libraries were sequenced on an Illumina NextSeq 500, generating ∼75 million reads per library. Read alignment was done using a custom pipeline with Bowtie 2 ^40, 41^.

## Supporting information

Method of chemical synthesis

## ACKNOWLEDGMENTS

We thank Justin Brache for preparation of caseum surrogate, and all members of the Sherman Lab for helpful discussions. This work was supported by a sub-award from NIH P30AI168034 to HE, and by NIH U19 AI 1625598 to DRS.

## Notes

### Competing Interest Statement

The authors have declared no competing interest.

## References

1. Dartois VA, Rubin EJ. Anti-tuberculosis treatment strategies and drug development: challenges and priorities. Nat Rev Microbiol 2022; 20: 685–701.

2. Zong Z, Huo F, Shi J et al. Relapse Versus Reinfection of Recurrent Tuberculosis Patients in a National Tuberculosis Specialized Hospital in Beijing, China. Front Microbiol 2018; 9: 1858.

3. Ragonnet R, Trauer JM, Denholm JT et al. High rates of multidrug-resistant and rifampicin-resistant tuberculosis among re-treatment cases: where do they come from? BMC Infect Dis 2017; 17: 36.

4. Cox H, Kebede Y, Allamuratova S et al. Tuberculosis recurrence and mortality after successful treatment: impact of drug resistance. PLoS Med 2006; 3: e384.

5. Jones RM, Adams KN, Eldesouky HE et al. The evolving biology of Mycobacterium tuberculosis drug resistance. Front Cell Infect Microbiol 2022; 12: 1027394.

6. Sebastian J, Swaminath S, Nair RR et al. De Novo Emergence of Genetically Resistant Mutants of Mycobacterium tuberculosis from the Persistence Phase Cells Formed against Antituberculosis Drugs In Vitro. Antimicrob Agents Chemother 2017; 61.

7. Cohen NR, Lobritz MA, Collins JJ. Microbial persistence and the road to drug resistance. Cell Host Microbe 2013; 13: 632–42.

8. Windels EM, Michiels JE, Fauvart M et al. Bacterial persistence promotes the evolution of antibiotic resistance by increasing survival and mutation rates. ISME J 2019; 13: 1239–51.

9. Vilchèze C, Jacobs WR. The Isoniazid Paradigm of Killing, Resistance, and Persistence in Mycobacterium tuberculosis. J Mol Biol 2019; 431: 3450–61.

10. Parvathaneni V, Kulkarni NS, Muth A et al. Drug repurposing: a promising tool to accelerate the drug discovery process. Drug Discov Today 2019; 24: 2076–85.

11. Farha MA, Brown ED. Drug repurposing for antimicrobial discovery. Nat Microbiol 2019; 4: 565–77.

12. Balaban NQ, Helaine S, Lewis K et al. Publisher Correction: Definitions and guidelines for research on antibiotic persistence. Nat Rev Microbiol 2019; 17: 460.

13. Grant SS, Hung DT. Persistent bacterial infections, antibiotic tolerance, and the oxidative stress response. Virulence 2013; 4: 273–83.

14. Sarathy JP, Dartois V. Caseum: a Niche for Mycobacterium tuberculosis Drug-Tolerant Persisters. Clin Microbiol Rev 2020; 33.

15. Joshi H, Kandari D, Bhatnagar R. Insights into the molecular determinants involved in. Virulence 2021; 12: 2721–49.

16. Saeed DK, Shakoor S, Razzak SA et al. Variants associated with Bedaquiline (BDQ) resistance identified in Rv0678 and eglux pump genes in Mycobacterium tuberculosis isolates from BDQ naïve TB patients in Pakistan. BMC Microbiol 2022; 22: 62.

17. Sonnenkalb L, Carter JJ, Spitaleri A et al. Bedaquiline and clofazimine resistance in Mycobacterium tuberculosis: an in-vitro and in-silico data analysis. Lancet Microbe 2023; 4: e358–e68.

18. Meikle V, Zhang L, Niederweis M. Intricate link between siderophore secretion and drug eglux in. Antimicrob Agents Chemother 2023; 67: e0162922.

19. Xu J, Li D, Shi J et al. Bedquiline Resistance Mutations: Correlations with Drug Exposures and Impact on the Proteome in M. tuberculosis. Antimicrob Agents Chemother 2023; 67: e0153222.

20. Baek SH, Li AH, Sassetti CM. Metabolic regulation of mycobacterial growth and antibiotic sensitivity. PLoS Biol 2011; 9: e1001065.

21. Pontes MH, Groisman EA. Slow growth determines nonheritable antibiotic resistance in. Sci Signal 2019; 12.

22. Stokes JM, Lopatkin AJ, Lobritz MA et al. Bacterial Metabolism and Antibiotic Egicacy. Cell Metab 2019; 30: 251–9.

23. Baer CE, Rubin EJ, Sassetti CM. New insights into TB physiology suggest untapped therapeutic opportunities. Immunol Rev 2015; 264: 327–43.

24. Connolly LE, Edelstein PH, Ramakrishnan L. Why is long-term therapy required to cure tuberculosis? PLoS Med 2007; 4: e120.

25. Yamamoto K, Nakata N, Mukai T et al. Coexpression of MmpS5 and MmpL5 Contributes to Both Eglux Transporter MmpL5 Trimerization and Drug Resistance in Mycobacterium tuberculosis. mSphere 2021; 6.

26. Shirley M. Netupitant/Palonosetron: A Review in Chemotherapy-Induced Nausea and Vomiting. Drugs 2021; 81: 1331–42.

27. Joseph DB. Pharmacology/toxicology NDA Review and evaluation of Akynzeo (https://www.accessdata.fda.gov/drugsatfda_docs/nda/2014/205718Orig1s000PharmR.pdf). Pharmacology Reviews. FDA-Center for drug evaluation and research, 2014.

28. Peterson EJR, Ma S, Sherman DR et al. Network analysis identifies Rv0324 and Rv0880 as regulators of bedaquiline tolerance in Mycobacterium tuberculosis. Nat Microbiol 2016; 1: 16078.

29. CLSI. Susceptibility Testing of Mycobacteria, Nocardia, and Other Aerobic Actinomycetes, 3rd ed. CLSI standard M24. Wayne, PA: Clinical and Laboratory Standards Institute., 2018.

30. Abou Mourad Ferreira M, Candeias Dos Santos L, Schmidt Castellani LG et al. Application of BactTiter-Glo ATP bioluminescence assay for Mycobacterium tuberculosis detection. Diagn Microbiol Infect Dis 2024; 109: 116275.

31. Omollo C, Singh V, Kigondu E et al. Developing synergistic drug combinations to restore antibiotic sensitivity in drug-resistant. Antimicrob Agents Chemother 2023; 65.

32. Bruhn DF, Scherman MS, Liu J et al. In vitro and in vivo Evaluation of Synergism between Anti-Tubercular Spectinamides and Non-Classical Tuberculosis Antibiotics. Sci Rep 2015; 5: 13985.

33. Chen P, Gearhart J, Protopopova M et al. Synergistic interactions of SQ109, a new ethylene diamine, with front-line antitubercular drugs in vitro. J Antimicrob Chemother 2006; 58: 332–7.

34. Peterson EJR, Brooks AN, Reiss DJ et al. MtrA modulates Mycobacterium tuberculosis cell division in host microenvironments to mediate intrinsic resistance and drug tolerance. Cell Rep 2023; 42: 112875.

35. Ferro BE, van Ingen J, Wattenberg M et al. Time-kill kinetics of antibiotics active against rapidly growing mycobacteria. J Antimicrob Chemother 2015; 70: 811–7.

36. Sarathy JP, Xie M, Jones RM et al. A Novel Tool to Identify Bactericidal Compounds against Vulnerable Targets in Drug-Tolerant M. tuberculosis found in Caseum. mBio 2023; 14: e0059823.

37. Rustad TR, Minch KJ, Ma S et al. Mapping and manipulating the Mycobacterium tuberculosis transcriptome using a transcription factor overexpression-derived regulatory network. Genome Biol 2014; 15: 502.

38. Ma S, Morrison R, Hobbs SJ et al. Transcriptional regulator-induced phenotype screen reveals drug potentiators in Mycobacterium tuberculosis. Nat Microbiol 2021; 6: 44–50.

39. Ma S, Jaipalli S, Larkins-Ford J et al. Transcriptomic Signatures Predict Regulators of Drug Synergy and Clinical Regimen Egicacy against Tuberculosis. mBio 2019; 10.

40. Langmead B, Salzberg SL. Fast gapped-read alignment with Bowtie 2. Nat Methods 2012; 9: 357–9.

41. Li H, Handsaker B, Wysoker A et al. The Sequence Alignment/Map format and SAMtools. Bioinformatics 2009; 25: 2078–9.

