## Supplemental Table 1, Supplemental Figure 1, Supplemental Figure 2, and Supplemental Figure 3 for "Netupitant Exhibits Potent Activity on *Mycobacterium tuberculosis* Persisters"

**Supplementary Table 1**. Minimum inhibitory concentration of netupitant (NTP) and the anti-TB drugs; isoniazid (INH), rifampicin (RIF), ethambutol (EMB), moxifloxacin (MOX), amikacin (AMK), and bedaquiline (BDQ).

| Name | MIC (μg/ml) |
|---|---|
| NTP | 16 |
| INH | 0.03125 |
| RIF | 0.004 |
| EMB | 1 |
| MOX | 0.0625 |
| AMK | 0.25 |
| BDQ | 0.125 |

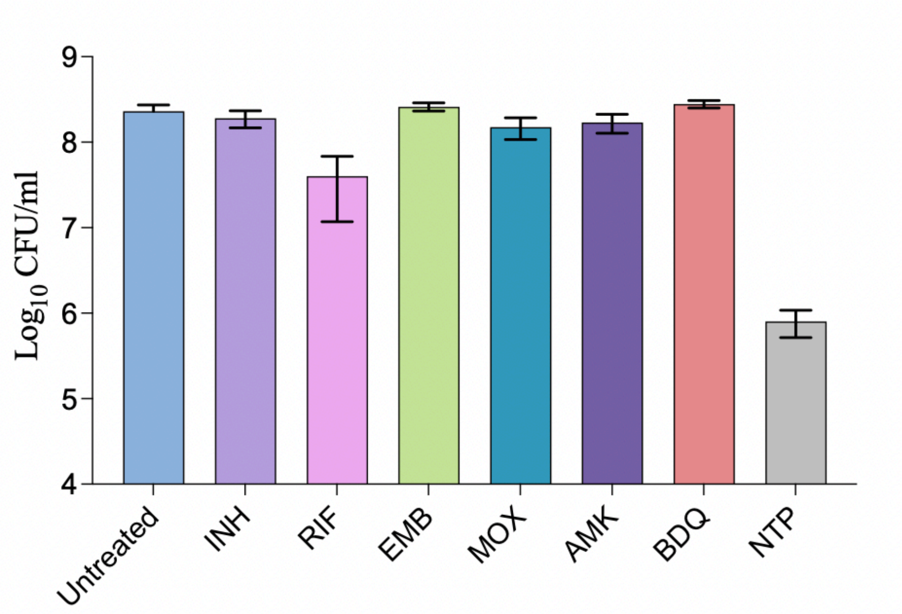


**Supplementary Figure 1: Impact of NTP and various anti-TB drugs on the viability of *M. tuberculosis* under hypoxic conditions.** *M. tuberculosis* H37Rv cultures were exposed to NTP (at 2x MIC) or a high dose (equivalent to 10x MIC) of isoniazid (INH), rifampicin (RIF), ethambutol (EMB), moxifloxacin (MOX), amikacin (AMK), or bedaquiline (BDQ). Cultures were then incubated under hypoxic conditions at 37°C for 7 days. Colony-forming units (CFU) were determined by plating on 7H10 agar and counted 21 days post-incubation at 37°C.

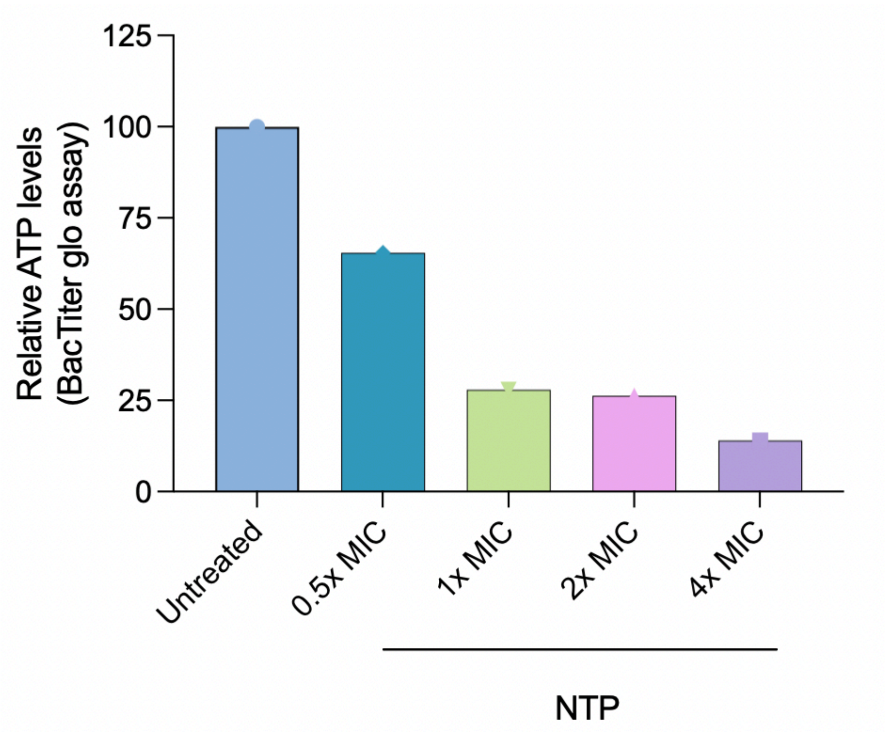


**Supplementary Figure 2: Effect of NTP on cellular ATP levels.** *M. tuberculosis* H37Rv cells were treated with different concentrations of NTP and incubated at 37 $^oC$ for 7 days. ATP levels were determined by the BacTiter glo assay and data were presented as the means $\pm$ standard deviations of the treated cells relative to the untreated control.

a

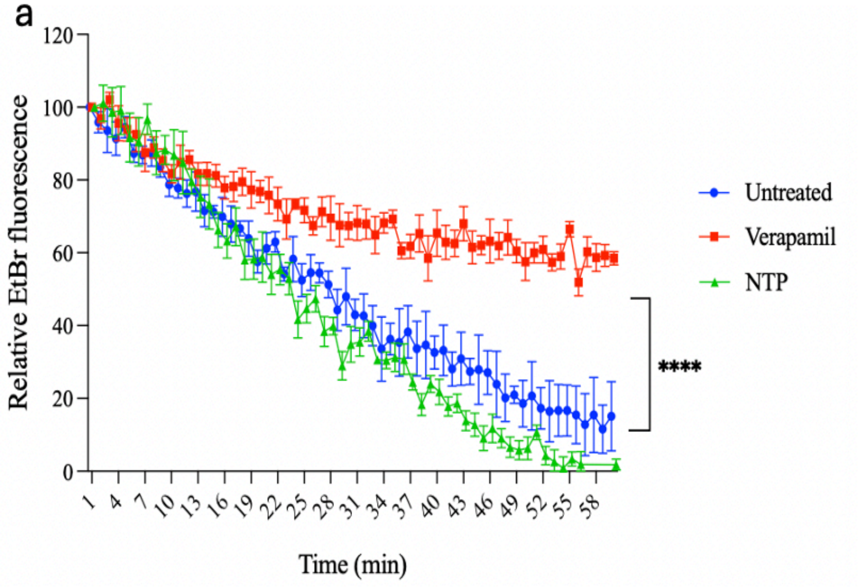


b

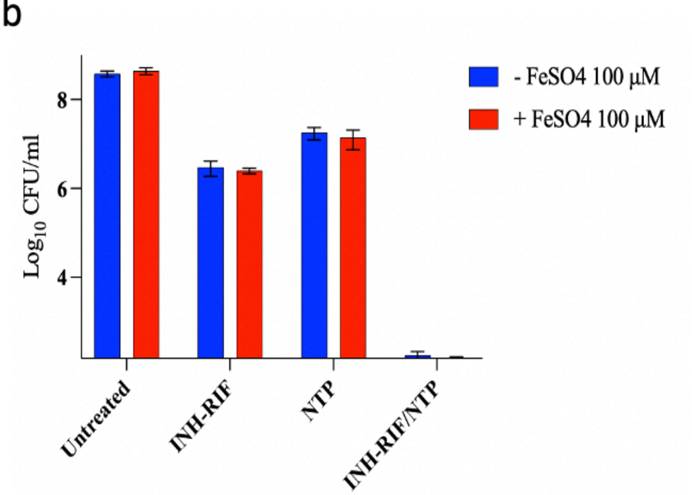


**Supplementary Figure 3: Activity of NTP on Rv0678 related biological functions.** (a) Effect of NTP (0.25x MIC) on ethidium bromide (EtBr) efflux in *M. tuberculosis* H37Rv cells, with verapamil (0.25x MIC) used as a positive control. (b) Impact of iron supplementation ($FeSO_4$ at 100 µg/ml) on *M. tuberculosis* H37Rv after 7 days of treatment with NTP (2x MIC) in the presence or absence of INH-RIF (100x MIC).
