## Supplementary material for "Netupitant Exhibits Potent Activity on *Mycobacterium tuberculosis* Persisters": Method of chemical synthesis

### Materials and Methods

#### Chemistry

#### Organic Synthesis

Commercially obtained reagents were used without further purification. Solvents were purchased as commercial anhydrous grade and used directly without additional treatment. Flash column chromatography was conducted using a CombiFlash instrument equipped with Flash Pure Buchi columns. Proton nuclear magnetic resonance ($^1H$ NMR) spectra were acquired on a 400 MHz Bruker spectrometer. High-resolution ESI-MS spectra were recorded using a Thermo Fisher Orbitrap Velos with an autosampler. Low-resolution mass spectrometry was performed via liquid chromatography–mass spectrometry (LC/MS) on a Waters Acquity UPLC system, utilizing either atmospheric pressure chemical ionization (APCI) or electrospray ionization (ESI) as needed.

The syntheses NTP, NTP-44, NTP-45, NTP-47, NTP-51, NTP-53 and NTP-59 are illustrated in schemes 1 and 2 shown below.

**Scheme 1.**

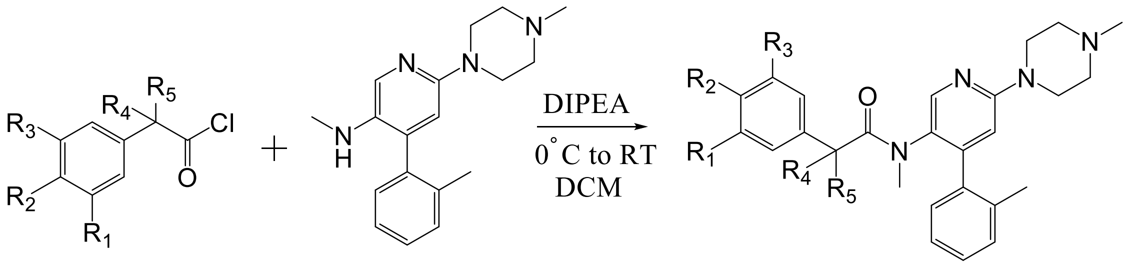

**1** **2** **3**

**NTP** $R_1$ and $R_3 = CF_3$, $R_2$ = H, $R_4$ and $R_5 = CH_3$

**NTP-44** $R_1$ and $R_3$ = H, $R_2$ = OMe, $R_4$ and $R_5 = CH_3$

**NTP-45** $R_{1,}$ $R_2$ and $R_3$ = H, $R_4$ and $R_5 = CH_3$

**NTP-47** $R_1$ and $R_3$ = H, $R_2$ = Cl, $R_4$ and $R_5 = CH_3$

**NTP-53** $R_1$ and $R_3 = CF_3$, $R_2$ = H, $R_4$ and $R_5$ = H

**Scheme 2**

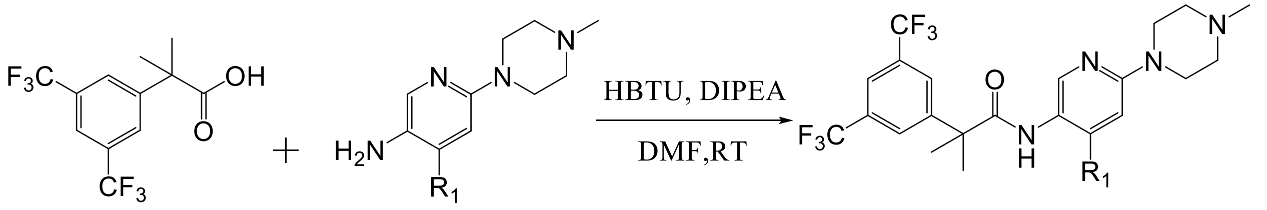

**4** **5**

**5a** = $R_1$ = *o*-tolyl

**5b** = $R_1$ = H

**NTP-51** $R_1$ = *o*-tolyl

**NTP-59** $R_1$ = H

**Experimental Procedure:**

**Syntheses of Netupitant and analogs**

**Synthesis of 2-(3,5-bis(trifluoromethyl)phenyl)-N,2-dimethyl-N-(6-(4-methylpiperazin-1-yl)-4-(o-tolyl)pyridin-3-yl)propenamide (Netupitant (NTP))**

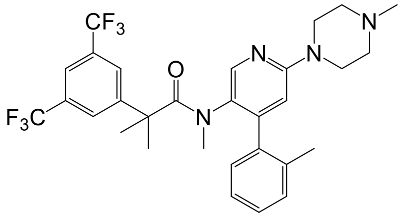

To a solution of N-methyl-6-(4-methylpiperazin-1-yl)-4-(o-tolyl)pyridin-3-amine (2, 1 mmol) in dichloromethane (DCM), the corresponding phenyl-2-methylpropanoyl chloride (1, 1.1 mmol)

was added at 0°C. Subsequently, DIPEA (1.5 mmol) was added, and the reaction mixture was stirred overnight. The mixture was then diluted with DCM and water, followed by extraction with DCM (2 × 20 mL). The combined organic layers were washed with an aqueous 2N $NaHCO_3$ solution, chilled water, dried over $Na_2SO_4$, and concentrated under reduced pressure. The crude product was purified by Combi Flash using a normal-phase column and a 0-15% DCM gradient in methanol (MeOH). Both the silica gel and the column cartridge were pre-treated with $Et_3N$, yielding the final compound, NTP. (White solid, 62% yield): $^1$HNMR (400 MHz, $CD_3OD$) δ 7.97 - 7.91 (m, 2H), 7.78 (bs, 2H), 7.32 - 7.23 (m, 4H), 6.91 (s, 1H), 3.63 (bs, 4H), (3H, merged with $CD_3OD$), 2.64 (t, *J* = 4.8 Hz, 4H), 2.41 (s, 3H), 2.1 (bs, 3H), 1.50 - 1.3 (m, 6H); LC-MS (ESI); [M+H]: 579.3

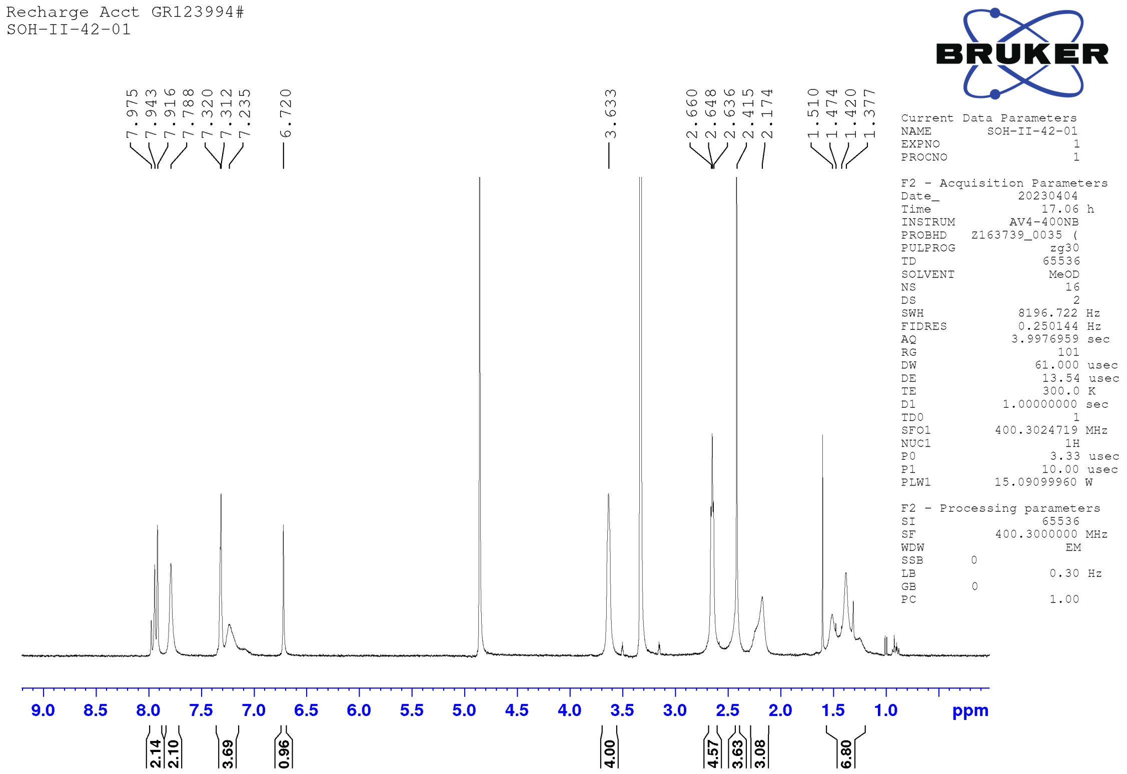

**Sample Report (continued):**

**Sample 3 SOH-II-42-01 23-Oct-2024 16:57:01 File: mohammed210_159-3-20241023-1641_COD**

1: MS ES+ :TIC Smooth (SG, 2x3) 2.6e+007

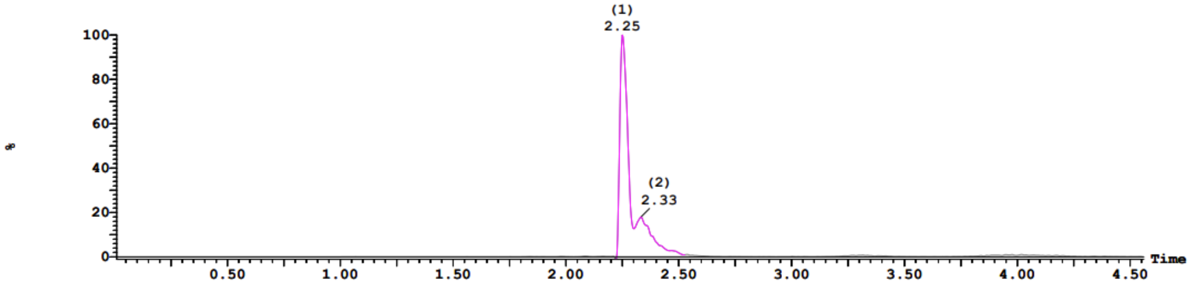


2: UV Detector: TAC: Wavelength Range: (210 - 498) 1.853e+1
Range: 1.853e+1

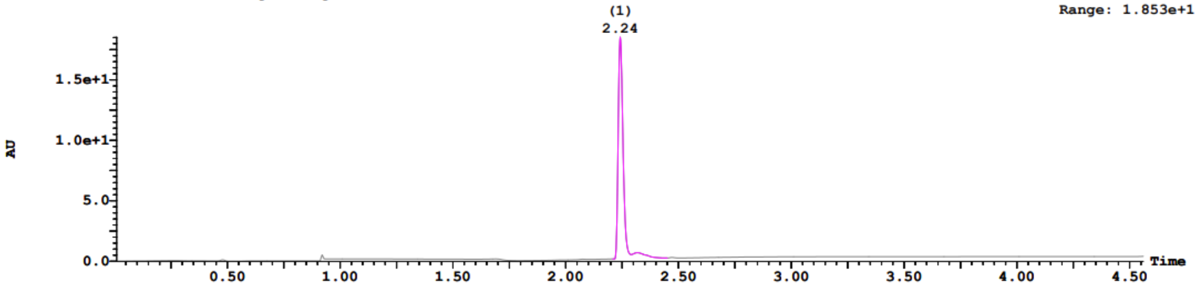


**Sample Report (continued):**

| Peak ID | Time |
|---|---|
| 1 | 2.25 |

1:(Time: 2.24) Combine (343:403-(313:342+404:433)) 1:MS ES+ 3.4e+006

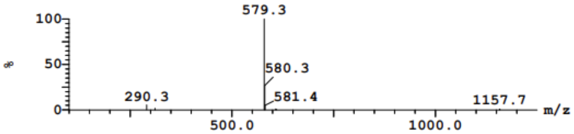


| Peak ID | Time |
|---|---|
| 1 | 2.25 |

1:(Time: 2.24) Combine (1340) 2:UV Detector 6.727e-1 AU

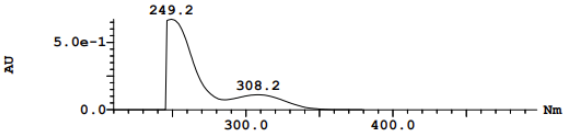


| Peak ID | Time |
|---|---|
| 2 | 2.33 |

2:(Time: 2.33) Combine (370:430-(340:369+431:460)) 1:MS ES+ 4.6e+006

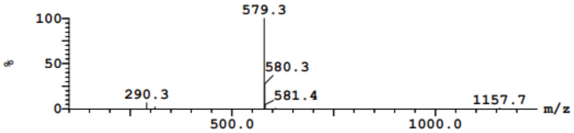


| Peak ID | Time |
|---|---|
| 2 | 2.33 |

2:(Time: 2.32) Combine (1386) 2:UV Detector 5.608e-2 AU

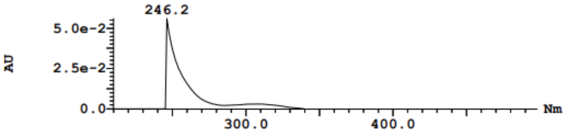


| Peak ID | Time |
|---|---|
| 3 | 2.35 |

3:(Time: 2.35) Combine (375:435-(345:374+436:465)) 1:MS ES+ 2.7e+006

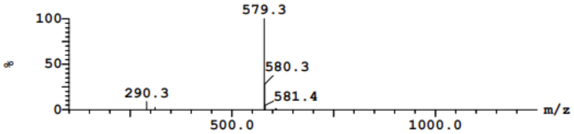


| Peak ID | Time |
|---|---|
| 3 | 2.35 |

3:(Time: 2.35) Combine (1406) 2:UV Detector 5.071e-2 AU

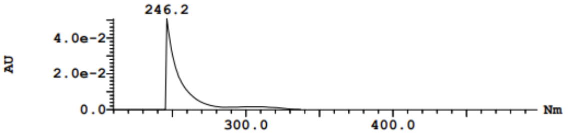

**Synthesis of 2-(4-methoxyphenyl)-N,2-dimethyl-N-(6-(4-methylpiperazin-1-yl)-4-(o-tolyl)pyridin-3-yl)propanamide (NTP-44)**

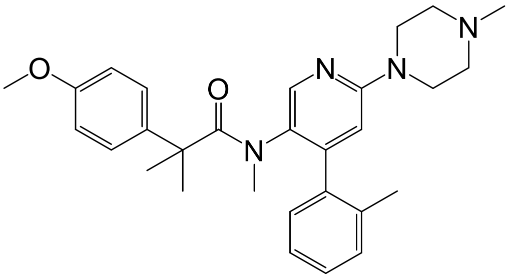


The synthesis of NTP-44 follows a procedure similar to that described for NTP. (White solid, 45% yield): [1]HNMR (400 MHz, $CDCl_3$) δ 8.0 (s, 1H), 7.34 - 7.24 (m, 4H), 6.82 (bs, 2H), 6.70 (distorted doublet, $J$ = 8.7 Hz, 2H), 6.48 (s, 1H), 3.77 (s, 3H), 3.58 (bs, 4H), 2.55 - 2.08 (m, 10H), 1.65 (bs, 3H), 1.49 - 1.25 (m, 6H); LC-MS (ESI); [M+H]: 473.4

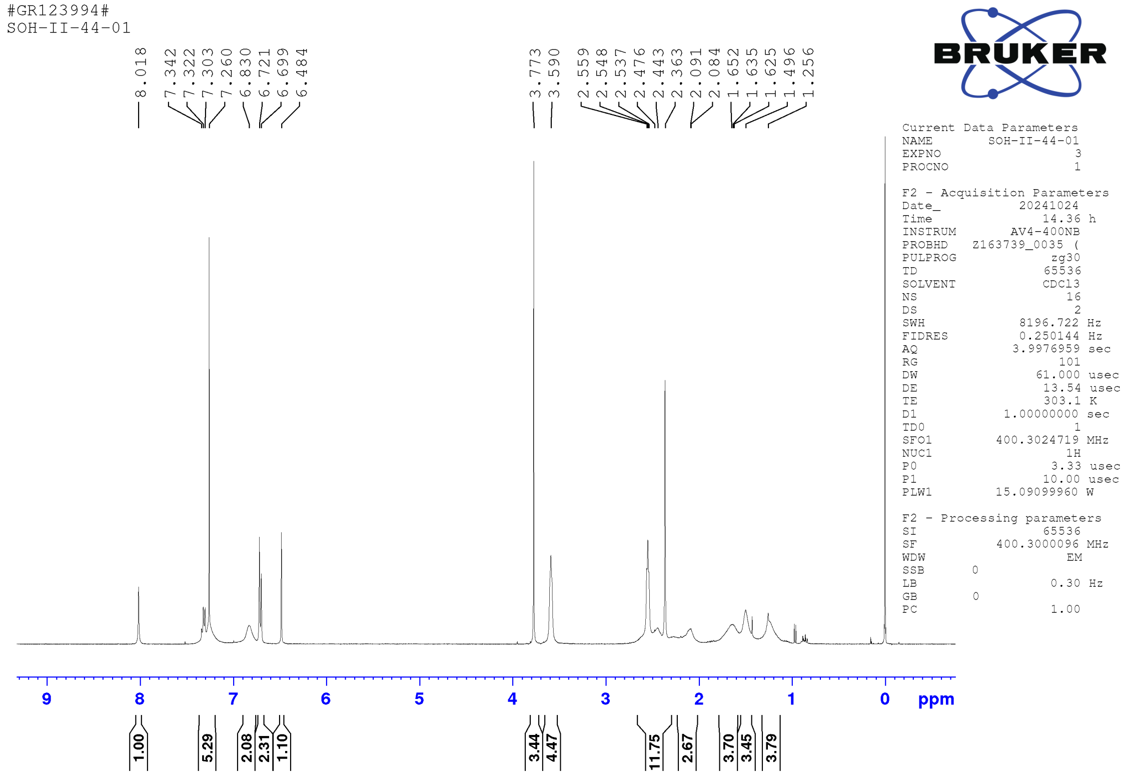

Vial:1:18 | ID:mohammed210_159-4-20241023-1641 | File:mohammed210_159-4-20241023-1641_COD
Date:23-Oct-2024 | Time:17:02:54 | Description:SOH-II-44-01
Method:C:\MassLynx\OpenLynx_Methods\ESI+_100-1250+PDA.olp | Instrument:ACQ-QDA#KBD6021
MS Method:OA_ESI+_Default | Inlet Method:OA_Default | Detectors:Waters Acquity PDA

Printed: Wed Oct 23 17:39:53 2024

**Sample Report (continued):**

**Sample 4 SOH-II-44-01 23-Oct-2024 17:02:54 File: mohammed210_159-4-20241023-1641_COD**

1: MS ES+ :TIC Smooth (SG, 2x3) 1.6e+007

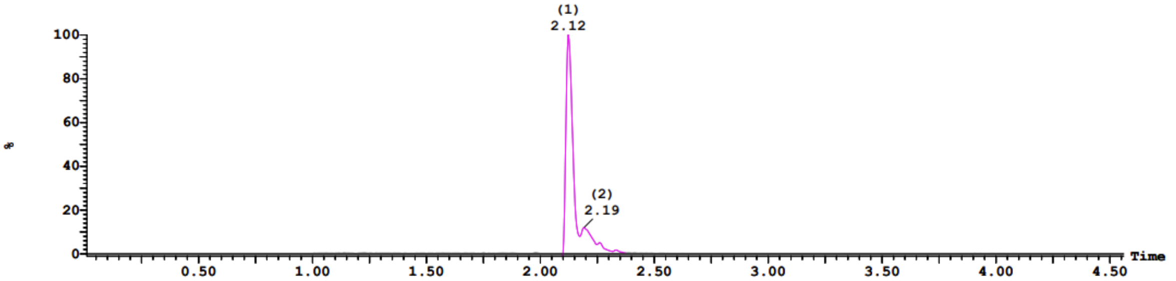


2: UV Detector: TAC: Wavelength Range: (210 - 498) 3.812
Range: 3.812

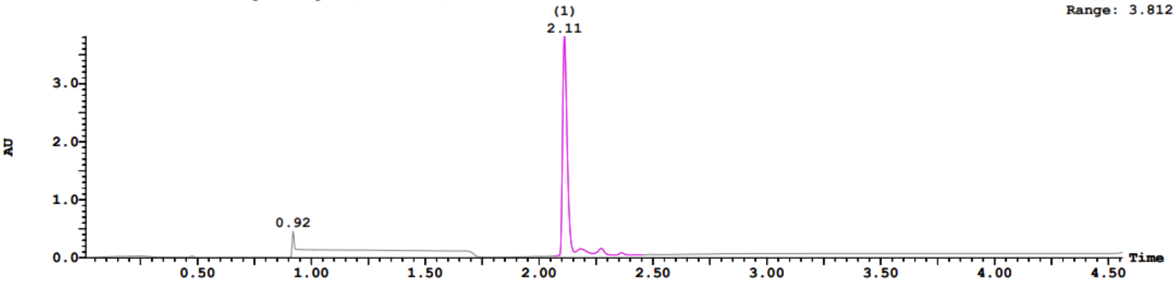


**Sample Report (continued):**

| Peak ID | Time |
|---|---|
| 1 | 2.12 |

1:(Time: 2.11) Combine (304:364-(274:303+365:394)) 1:MS ES+
1.2e+006

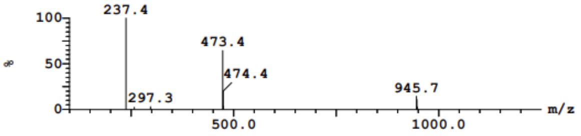


| Peak ID | Time |
|---|---|
| 1 | 2.12 |

1:(Time: 2.11) Combine (1262) 2:UV Detector
1.857e-1 AU

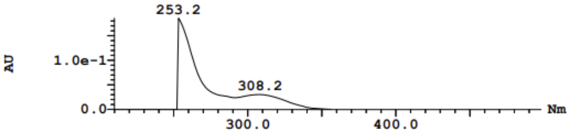


| Peak ID | Time |
|---|---|
| 2 | 2.19 |

2:(Time: 2.20) Combine (331:391-(301:330+392:421)) 1:MS ES+
1.4e+006

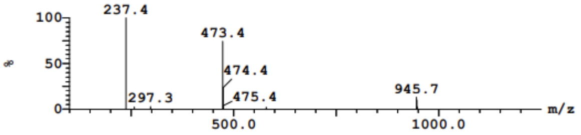


| Peak ID | Time |
|---|---|
| 2 | 2.19 |

2:(Time: 2.18) Combine (1305) 2:UV Detector
1.358e-2 AU

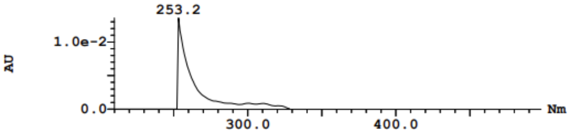


| Peak ID | Time |
|---|---|
| 3 | 2.26 |

3:(Time: 2.26) Combine (348:408-(318:347+409:438)) 1:MS ES+
3.2e+004

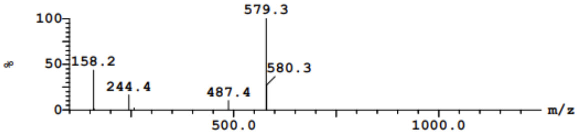


| Peak ID | Time |
|---|---|
| 3 | 2.26 |

3:(Time: 2.27) Combine (1359) 2:UV Detector
1.45e-2 AU

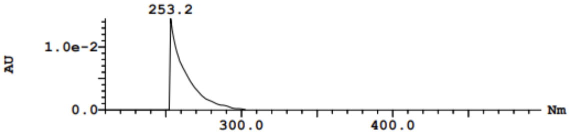

**Synthesis of N,2-dimethyl-N-(6-(4-methylpiperazin-1-yl)-4-(o-tolyl)pyridin-3-yl)-2-phenylpropanamide (NTP-45)**

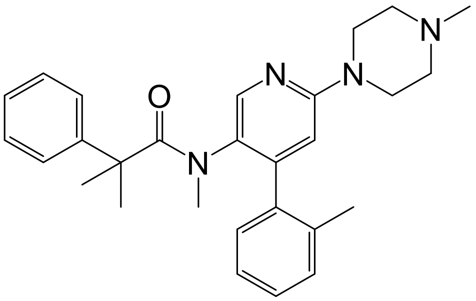


The synthesis of NTP-45 follows a procedure similar to that described for NTP (White solid, 47% yield): [1]HNMR (400 MHz, $CDCl_3$) δ 8.05 (s, 1H), 7.37 - 7.14 (m, 8H), 6.96 (bs, 1H), 6.51 (s, 1H), 3.63 (bs, 4H), 2.59 - 2.10 (m, 13H), 1.55 - 1.29 (m, 6H); LC-MS (ESI); [M+H]: 443.4

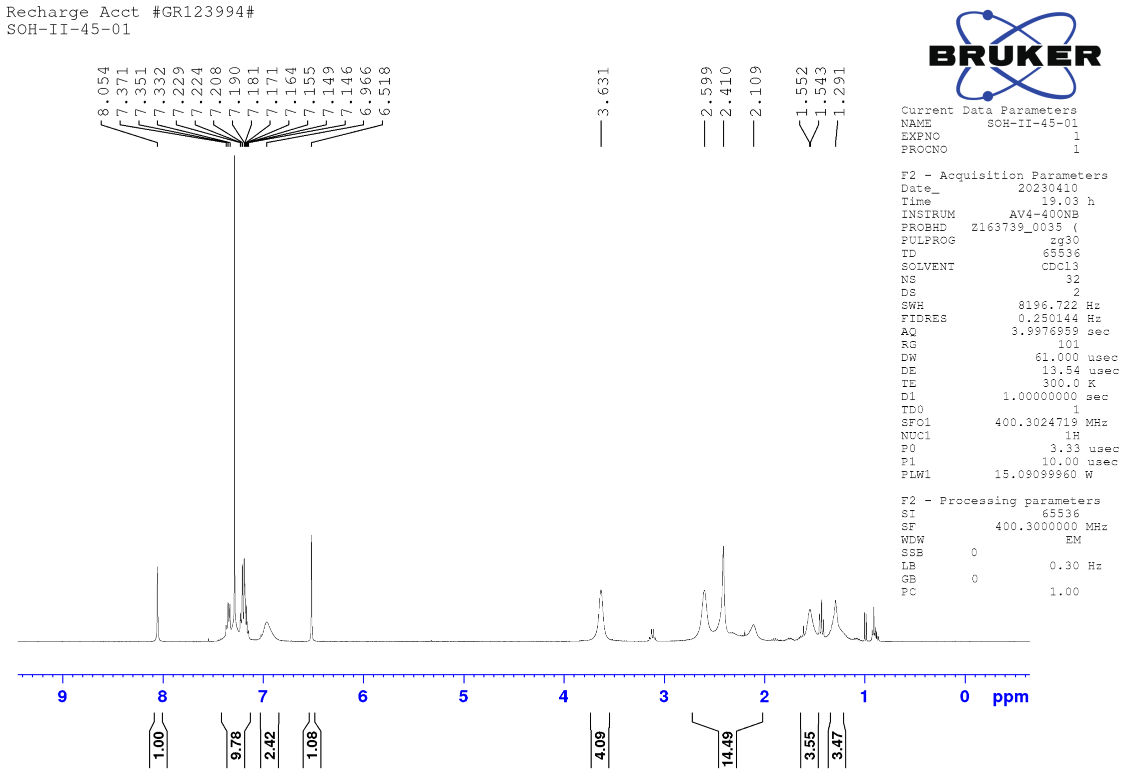

Vial:1:19
Date:23-Oct-2024
Method:C:\MassLynx\OpenLynx_Methods\ESI+_100-1250+PDA.olp
MS Method:OA_ESI+_Default
ID:mohammed210_159-5-20241023-1641
Time:17:08:48
Inlet Method:OA_Default
File:mohammed210_159-5-20241023-1641_COD
Description:SOH-II-45-01
Instrument:ACQ-QDA#KBD6021
Detectors:Waters Acquity PDA

Printed: Wed Oct 23 17:39:53 2024

**Sample Report (continued):**

**Sample 5 SOH-II-45-01 23-Oct-2024 17:08:48 File: mohammed210_159-5-20241023-1641_COD**

1: MS ES+ :TIC Smooth (SG, 2x3) 2.5e+007

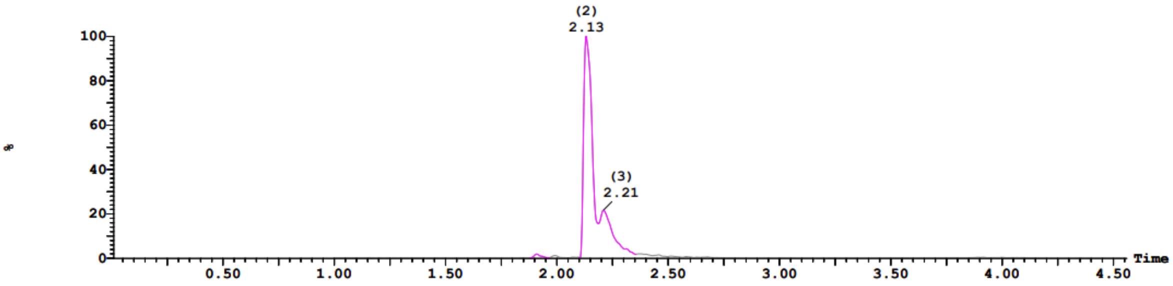


2: UV Detector: TAC: Wavelength Range: (210 - 498) 1.722e+1 Range: 1.722e+1

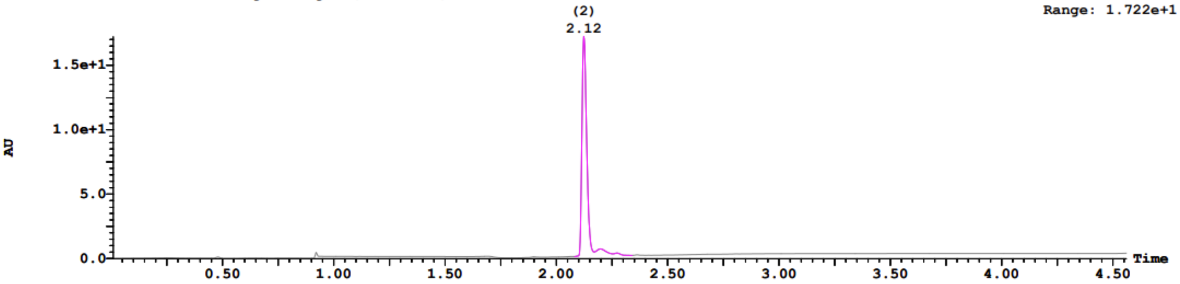


**Sample Report (continued):**

| Peak ID | Time |
|---|---|
| 1 | 1.91 |

1:(Time: 1.91) Combine (243:303-(213:242+304:333)) 1:MS ES+ 2.3e+004

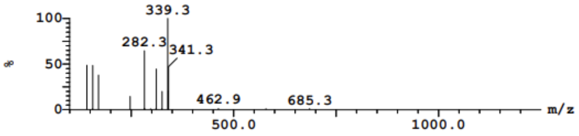


| Peak ID | Time |
|---|---|
| 2 | 2.13 |

2:(Time: 2.12) Combine (307:367-(277:306+368:397)) 1:MS ES+ 2.0e+006

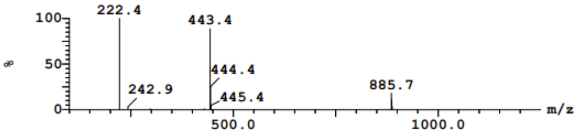


| Peak ID | Time |
|---|---|
| 2 | 2.13 |

2:(Time: 2.12) Combine (1269) 2:UV Detector 6.651e-1 AU

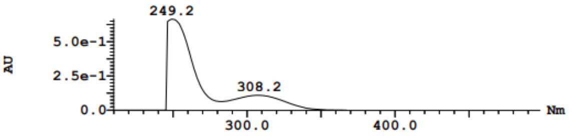


| Peak ID | Time |
|---|---|
| 3 | 2.21 |

3:(Time: 2.21) Combine (333:393-(303:332+394:423)) 1:MS ES+ 2.6e+006

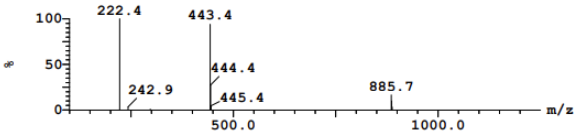


| Peak ID | Time |
|---|---|
| 3 | 2.21 |

3:(Time: 2.20) Combine (1314) 2:UV Detector 5.343e-2 AU

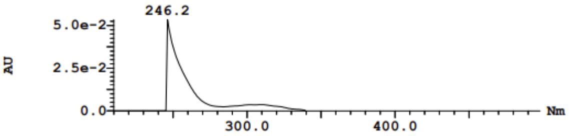


| Peak ID | Time |
|---|---|
| 4 | 2.23 |

4:(Time: 2.23) Combine (340:400-(310:339+401:430)) 1:MS ES+ 1.5e+006

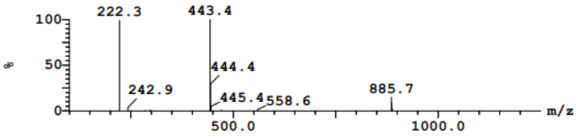

**Synthesis of 2-(4-chlorophenyl)-N,2-dimethyl-N-(6-(4-methylpiperazin-1-yl)-4-(o-tolyl)pyridin-3-yl)propanamide (NTP-47)**

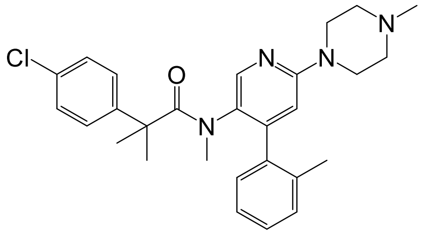


The synthesis of NTP-47 follows a procedure similar to that described for NTP (White solid, 62% yield): $^1$HNMR (400 MHz, $CDCl_3$) δ 8.0 (s, 1H), 7.36 - 7.24 (m, 5H), 7.12 (d, *J* = 8.5 Hz, 2H), 6.81 (bs, 1H), 6.48 (s, 1H), 3.58 (t, *J* = 4.7 Hz, 4H), 2.55 - 2.08 (m, 10H), 1.74 (bs, 3H), 1.52 - 1.25 (m, 6H); LC-MS (ESI); [M+H]: 477.3

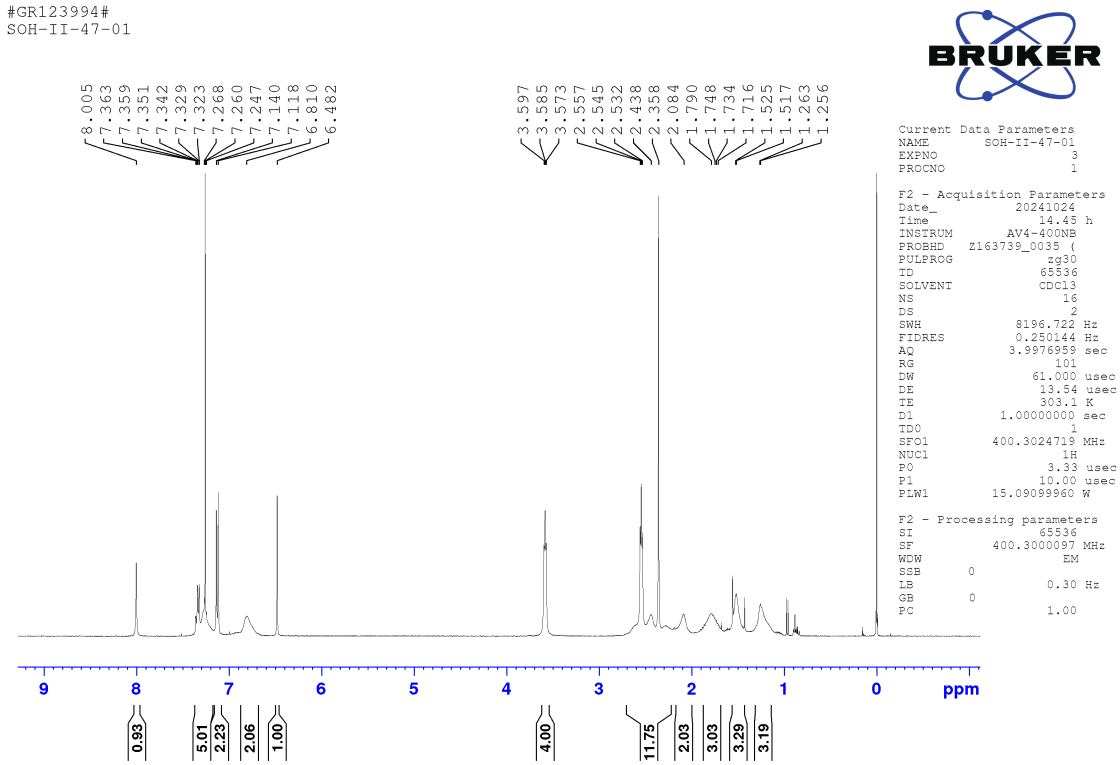

Vial:1:20
Date:23-Oct-2024
Method:C:\MassLynx\OpenLynx_Methods\ESI+_100-1250+PDA.olp
MS Method:OA_ESI+_Default

ID:mohammed210_159-6-20241023-1641
Time:17:14:43
Inlet Method:OA_Default

File:mohammed210_159-6-20241023-1641_COD
Description:SOH-II-47-01
Instrument:ACQ-QDA#KBD6021
Detectors:Waters Acquity PDA

Printed: Wed Oct 23 17:39:53 2024

**Sample Report (continued):**

**Sample 6 SOH-II-47-01 23-Oct-2024 17:14:43 File: mohammed210_159-6-20241023-1641_COD**

1: MS ES+ :TIC Smooth (SG, 2x3) 2.0e+007

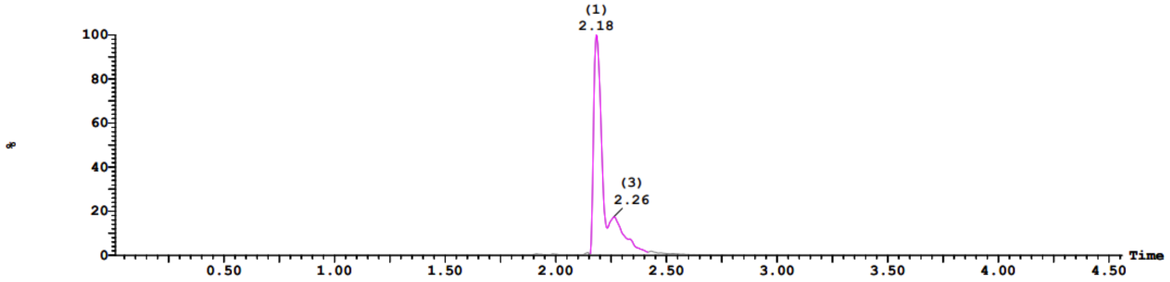


2: UV Detector: TAC: Wavelength Range: (210 - 498) 1.397e+1
Range: 1.397e+1

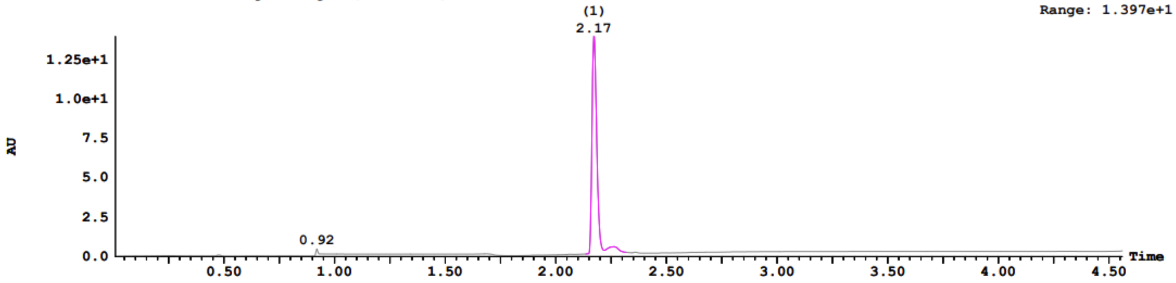


**Sample Report (continued):**

**Peak ID** 1 **Time** 2.18

1:(Time: 2.17) Combine (322:382-(292:321+383:412)) 1:MS ES+
1.7e+006

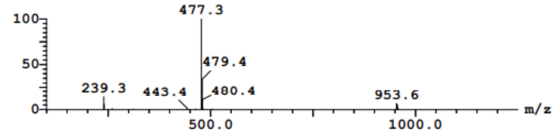


**Peak ID** 1 **Time** 2.18

1:(Time: 2.17) Combine (1298) 2:UV Detector
5.604e-1 AU

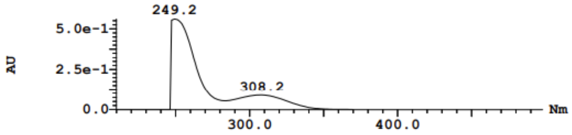


**Peak ID** 2 **Time** 2.26

2:(Time: 2.26) Combine (347:407-(317:346+408:437)) 1:MS ES+
2.4e+006

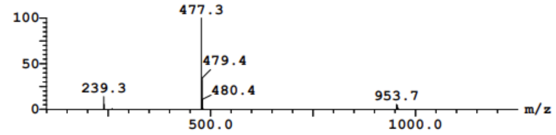


**Peak ID** 2 **Time** 2.26

2:(Time: 2.26) Combine (1348) 2:UV Detector
4.603e-2 AU

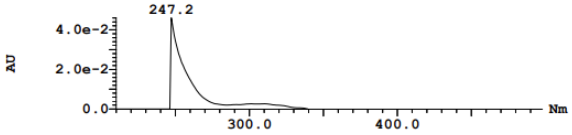


**Peak ID** 3 **Time** 2.26

3:(Time: 2.26) Combine (349:409-(319:348+410:439)) 1:MS ES+
2.4e+006

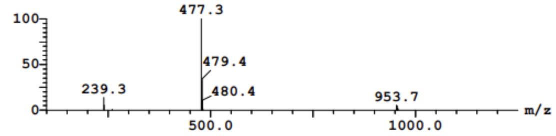


**Peak ID** 3 **Time** 2.26

3:(Time: 2.26) Combine (1353) 2:UV Detector
4.614e-2 AU

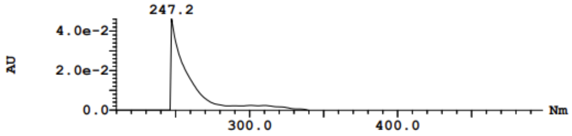

**Synthesis of 2-(3,5-bis(trifluoromethyl)phenyl)-N-(6-(4-methylpiperazin-1-yl)-4-(o-tolyl)pyridin-3-yl)acetamide (NTP-53)**

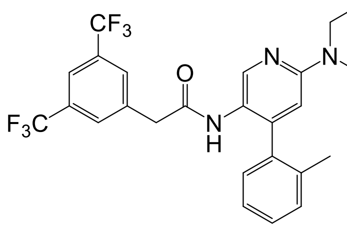


The synthesis of NTP-53 follows a procedure similar to that described for NTP (White solid, 30% yield): $^{1}$HNMR (400 MHz, $CDCl_3$) δ 8.05 (s, 1H), 7.76 (bs, 1H), 7.54 (bs, 2H), 7.36 - 7.30 (m, 2H), 7.22 (distorted triplet, *J* = 7.1 Hz, 1H), 6.95 (bs, 1H), 6.58 (bs, 1H), 3.66 (bs, 6H), 2.97 (bs, 3H), 2.58 (bs, 4H), 2.40 (s, 3H), 2.24 (s, 3H); LC-MS (ESI); [M+H]: 551.3

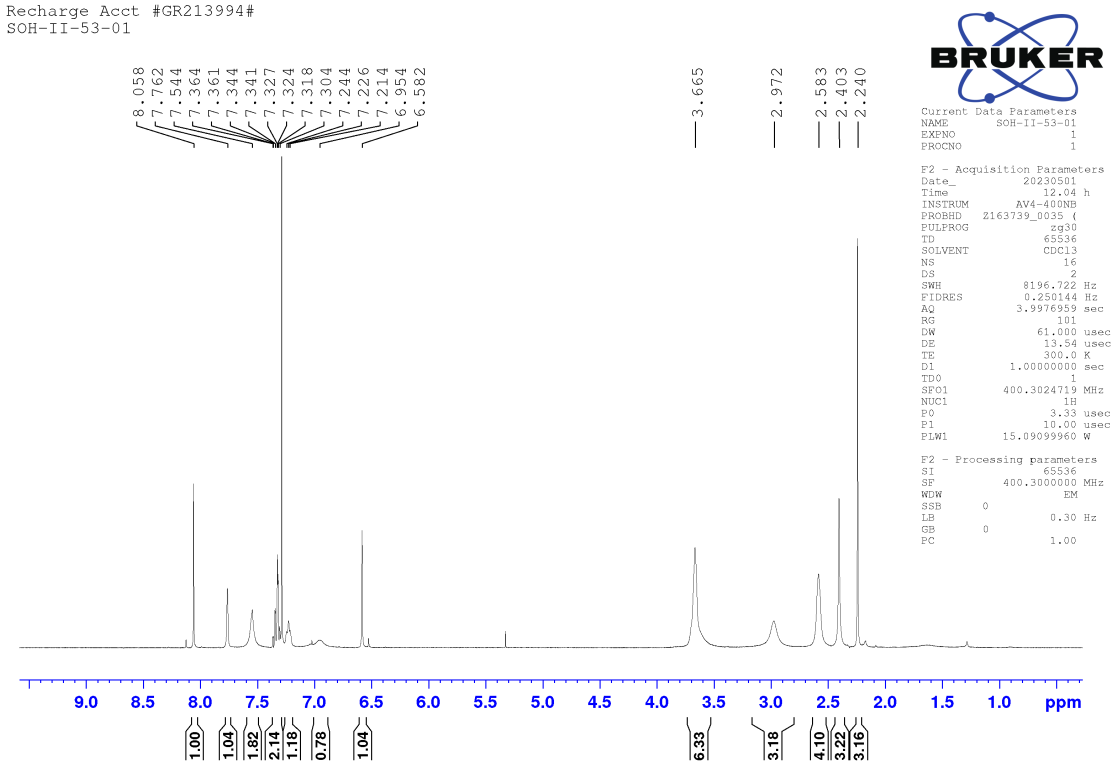

Vial:1:22 ID:mohammed210_159-8-20241023-1641 File:mohammed210_159-8-20241023-1641_COD
Date:23-Oct-2024 Time:17:26:36 Description:SOH-II-53-01
Method:C:\MassLynx\OpenLynx_Methods\ESI+_100-1250+PDA.olp Instrument:ACQ-QDA#KBD6021
MS Method:OA_ESI+_Default Inlet Method:OA_Default Detectors:Waters Acquity PDA

Printed: Wed Oct 23 17:39:53 2024

**Sample Report (continued):**

**Sample 8 SOH-II-53-01 23-Oct-2024 17:26:36 File: mohammed210_159-8-20241023-1641_COD**

1: MS ES+ :TIC Smooth (SG, 2x3) 2.7e+007

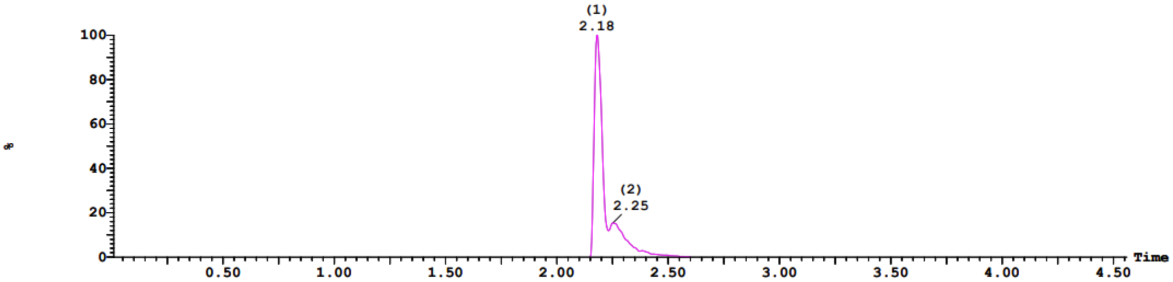


2: UV Detector: TAC: Wavelength Range: (210 - 498) 1.717e+1
Range: 1.717e+1

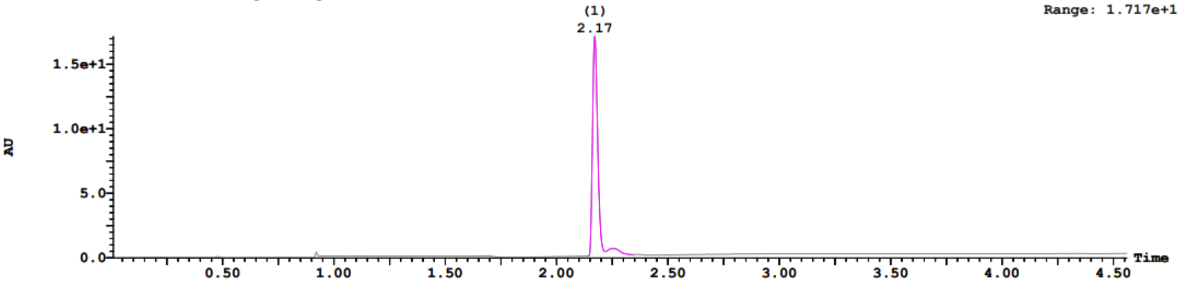


**Sample Report (continued):**

| Peak ID | Time |
|---|---|
| 1 | 2.18 |

1:(Time: 2.17) Combine (322:382-(292:321+383:412)) 1:MS ES+ 3.9e+006

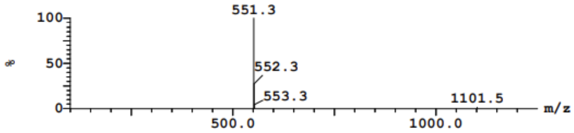


| Peak ID | Time |
|---|---|
| 1 | 2.18 |

1:(Time: 2.17) Combine (1298) 2:UV Detector 6.12e-1 AU

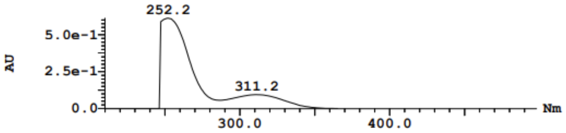


| Peak ID | Time |
|---|---|
| 2 | 2.25 |

2:(Time: 2.25) Combine (346:406-(316:345+407:436)) 1:MS ES+ 5.0e+006

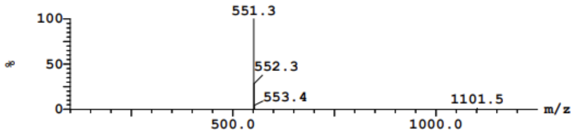


| Peak ID | Time |
|---|---|
| 2 | 2.25 |

2:(Time: 2.26) Combine (1349) 2:UV Detector 4.776e-2 AU

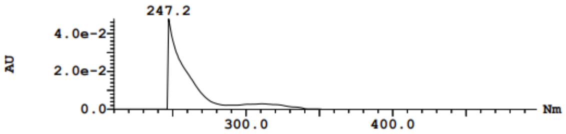


| Peak ID | Time |
|---|---|
| 3 | 2.28 |

3:(Time: 2.28) Combine (354:414-(324:353+415:444)) 1:MS ES+ 2.7e+006

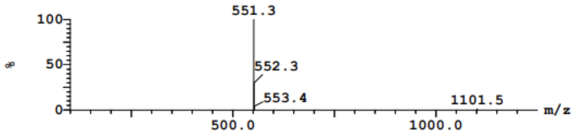


| Peak ID | Time |
|---|---|
| 4 | 2.31 |

4:(Time: 2.31) Combine (363:423-(333:362+424:453)) 1:MS ES+ 2.3e+004

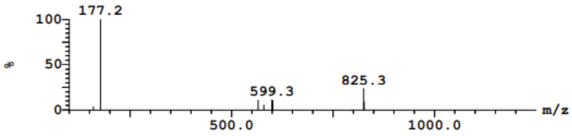

**Synthesis of 2-(3,5-bis(trifluoromethyl)phenyl)-2-methyl-N-(6-(4-methylpiperazin-1-yl)-4-(o-tolyl)pyridin-3-yl)propanamide (NTP-51)**

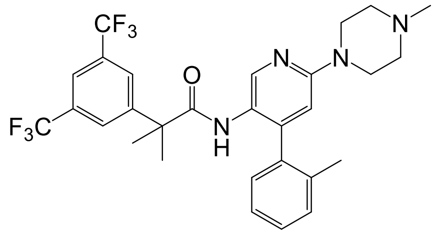


To a solution of 2-(3,5-bis(trifluoromethyl)phenyl)-2-methylpropanoic acid (4, 1.0 mmol) in DMF, HBTU (1.5 mmol) and DIPEA (1.1 mmol) were added sequentially, and the mixture was stirred for 15 minutes. Subsequently, 6-(4-methylpiperazin-1-yl)pyridin-3-amine (5, 1.1 mmol) was introduced to the reaction mixture and stirred overnight. The mixture was then diluted with EtOAc and water and extracted with EtOAc (3 × 20 mL). The combined organic layers were washed with 2N aqueous $NaHCO_3$, chilled water, dried over $Na_2SO_4$, and concentrated under reduced pressure. The crude product was purified by combi flash chromatography using a normal phase column with a 0-15% DCM in MeOH gradient. Both the silica gel and column cartridge were pre-treated with $Et_3N$ yielding NTP-51. (White solid, 35% yield): $^1$HNMR (400 MHz, $CDCl_3$) δ 9.07 (s, 1H), 7.84 (s, 1H), 7.70 - 7.68 (m, 1H), 7.57 (s, 2H), 7.20 (td, $J$ = 7.5 Hz, 1.2 Hz, 1H), 7.10 - 7.04 (m, 1H), 6.84 (dd, $J$ = 7.4 Hz, 1.0 Hz, 1H), 6.37 (s, 1H), 6.13 (s, 1H), 3.57 (t, $J$ = 4.3 Hz, 4H), 2.71 (t, $J$ = 4.8 Hz, 4H), 2.42 (s, 3H), 1.83 (s, 3H), 1.51 - 1.48 (m, 6H); LC-MS (ESI); [M+H]: 565.4

Recharge Acct #GR123994#
SOH-II-51-01

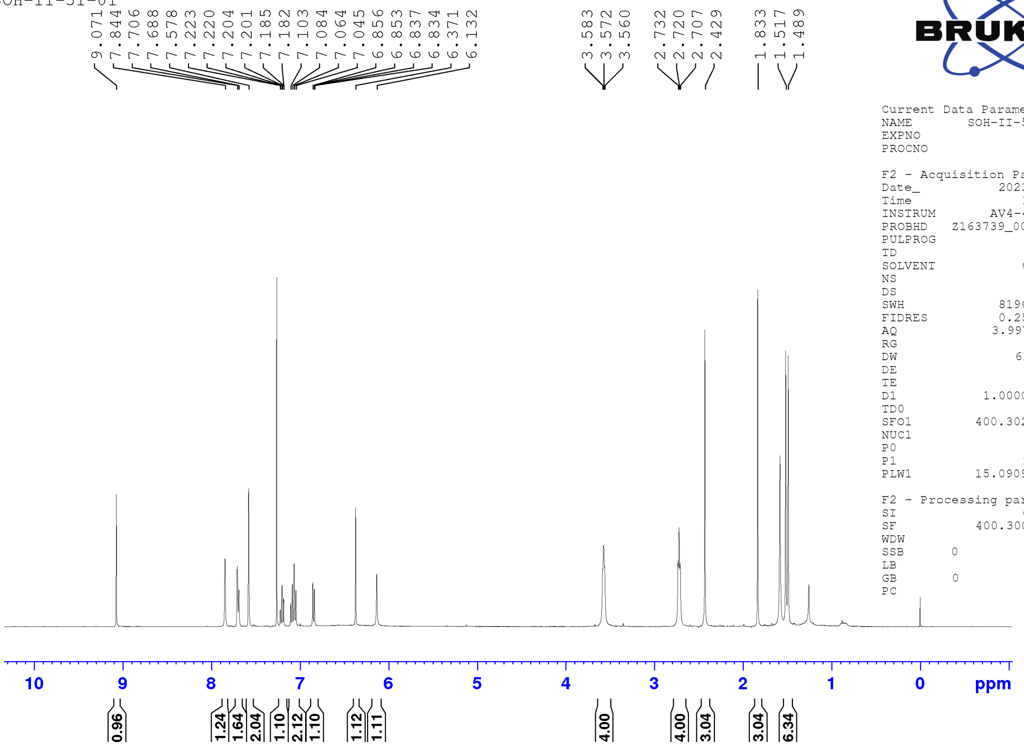


```
Current Data Parameters
NAME        SOH-II-51-01
EXPNO                  1
PROCNO                 1

F2 - Acquisition Parameters
Date_           20230510
Time               16.39 h
INSTRUM        AV4-400NB
PROBHD   Z163739_0035 (
PULPROG             zg30
TD                 65536
SOLVENT            CDCl3
NS                    16
DS                     2
SWH             8196.722 Hz
FIDRES          0.250144 Hz
AQ             3.9976959 sec
RG                   101
DW                61.000 usec
DE                 13.54 usec
TE                 300.0 K
D1            1.00000000 sec
TD0                    1
SFO1         400.3024719 MHz
NUC1                  1H
P0                  3.33 usec
P1                 10.00 usec
PLW1         15.09099960 W

F2 - Processing parameters
SI                 65536
SF           400.3000098 MHz
WDW                   EM
SSB      0
LB                  0.30 Hz
GB       0
PC                  1.00
```

**Openlynx Report – mohammed210**

Vial:1:21
Date:23-Oct-2024
Method:C:\MassLynx\OpenLynx_Methods\ESI+_100-1250+PDA.olp
MS Method:OA_ESI+_Default

ID:mohammed210_159-7-20241023-1641
Time:17:20:39
Inlet Method:OA_Default

File:mohammed210_159-7-20241023-1641_COD
Description:SOH-II-51-01
Instrument:ACQ-QDA#KBD6021
Detectors:Waters Acquity PDA

Printed: Wed Oct 23 17:39:53 2024

**Sample Report (continued):**

**Sample 7 SOH-II-51-01 23-Oct-2024 17:20:39 File: mohammed210_159-7-20241023-1641_COD**

1: MS ES+ :TIC Smooth (SG, 2x3) 1.4e+007

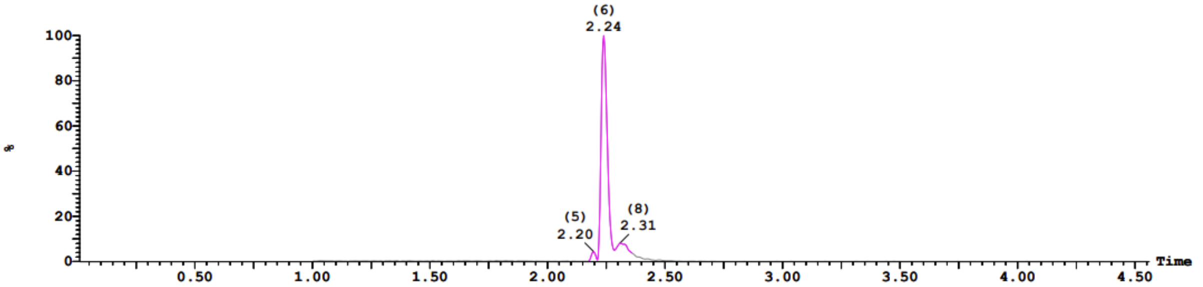


2: UV Detector: TAC: Wavelength Range: (210 - 498) 1.94
Range: 1.94

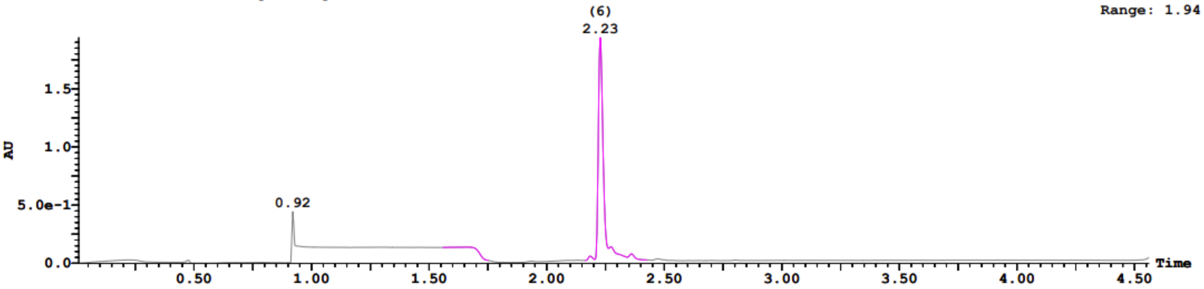


**Sample Report (continued):**

**Peak ID** **Time**
4 1.66
4: (Time: 1.67) Combine (171:231-(141:170+232:261)) 1:MS ES+
4.7e+003

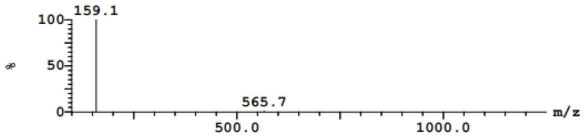


**Peak ID** **Time**
4 1.66
4: (Time: 1.67) Combine (996) 2:UV Detector
1.021e-3 AU

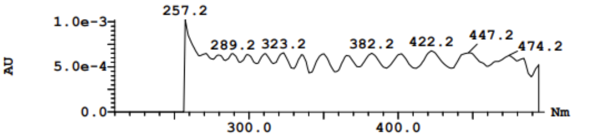


**Peak ID** **Time**
5 2.20
5: (Time: 2.20) Combine (329:389-(299:328+390:419)) 1:MS ES+
8.1e+005

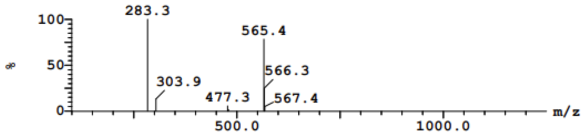


**Peak ID** **Time**
5 2.20
5: (Time: 2.18) Combine (1306) 2:UV Detector
5.193e-3 AU

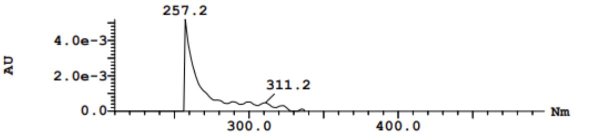


**Peak ID** **Time**
6 2.24
6: (Time: 2.24) Combine (342:402-(312:341+403:432)) 1:MS ES+
9.3e+005

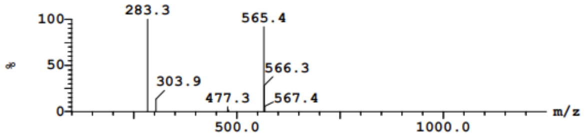


**Peak ID** **Time**
6 2.24
6: (Time: 2.23) Combine (1332) 2:UV Detector
6.528e-2 AU

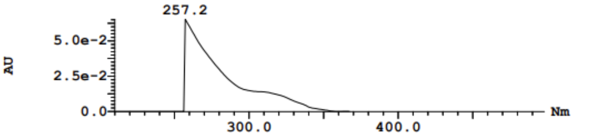

**Synthesis of 2-(3,5-bis(trifluoromethyl)phenyl)-2-methyl-N-(6-(4-methylpiperazin-1-yl)pyridin-3-yl)propanamide (NTP-59)**

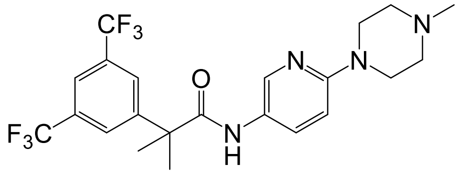


To a solution of 2-(3,5-bis(trifluoromethyl)phenyl)-2-methylpropanoic acid in DMF, HBTU (1.5 mmol) and DIPEA (1.1 mmol) were added sequentially, and the mixture was stirred for 15 minutes. Next, 6-(4-methylpiperazin-1-yl)pyridin-3-amine (1.1 mmol) was added, and the reaction was stirred overnight. The mixture was then purified by loading onto a combi flash chromatography with a normal-phase column using a 0-15% DCM in MeOH gradient. Both the silica gel and column cartridge were pre-treated with Et3N yielding NTP-59. (White solid, 56% yield): $^{1}$HNMR (400 MHz, DMSO-$d_6$) δ 7.33 (bs, 1H), 7.04 - 6.94 (m, 3H), 6.85 - 6.82 (m, 1H), 6.64 - 6.54 (m, 1H), 6.01 (t, $J$ = 8.9 Hz, 1H), 2.86 - 2.81 (bs, 4H), 2.44 (bs, 4H), 2.02 (s, 3H), 0.8 (s, 6H); LC-MS (ESI); [M+H]: 475.3

Recharge Acct #GR123994#
SOH-II-59-01

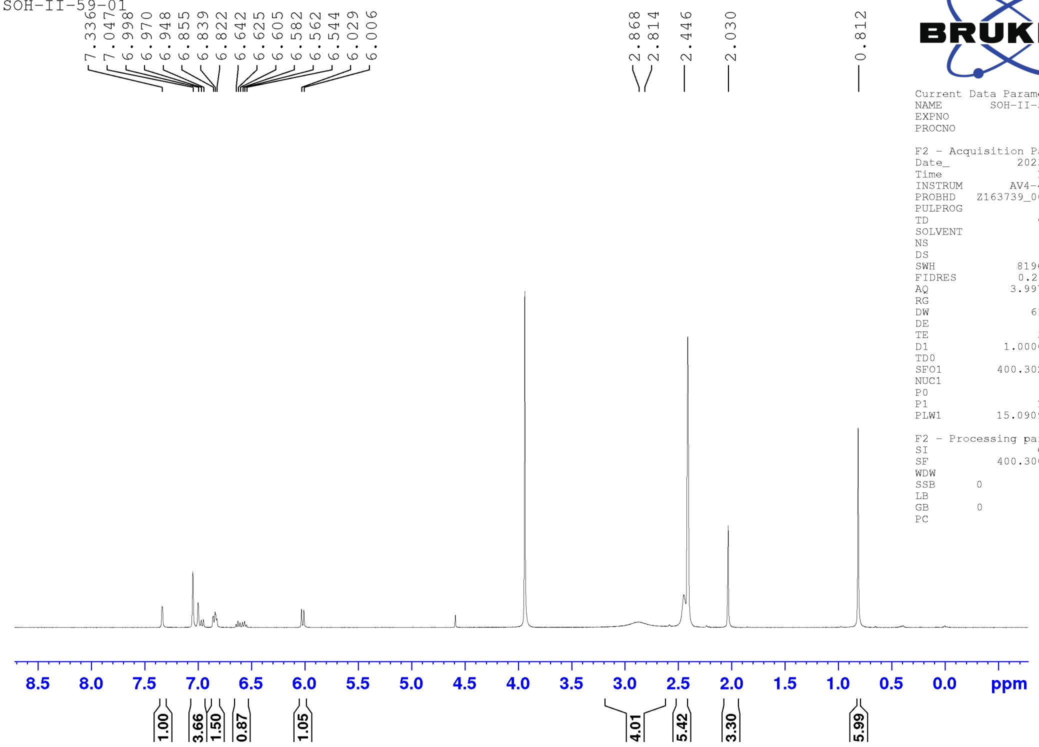


BRUKER

```
Current Data Parameters
NAME          SOH-II-59-01
EXPNO                    2
PROCNO                   1

F2 - Acquisition Parameters
Date_             20230508
Time                 16.17 h
INSTRUM          AV4-400NB
PROBHD    Z163739_0035 (
PULPROG               zg30
TD                   65536
SOLVENT               DMSO
NS                      16
DS                       2
SWH               8196.722 Hz
FIDRES            0.250144 Hz
AQ               3.9976959 sec
RG                     101
DW                  61.000 usec
DE                   13.54 usec
TE                   300.0 K
D1              1.00000000 sec
TD0                      1
SFO1           400.3024719 MHz
NUC1                    1H
P0                    3.33 usec
P1                   10.00 usec
PLW1           15.09099960 W

F2 - Processing parameters
SI                   65536
SF             400.3000443 MHz
WDW                     EM
SSB      0
LB                    0.30 Hz
GB       0
PC                    1.00
```

**Openlynx Report – mohammed210**

Vial:1:23
Date:23-Oct-2024
Method:C:\MassLynx\OpenLynx_Methods\ESI+_100-1250+PDA.olp
MS Method:OA_ESI+_Default

ID:mohammed210_159-9-20241023-1641
Time:17.32.32
Inlet Method:OA_Default

File:mohammed210_159-9-20241023-1641_COD
Description:SOH-II-59-01
Instrument:ACQ-QDA#KBD6021
Detectors:Waters Acquity PDA

Printed: Wed Oct 23 17:39:53 2024

**Sample Report (continued):**

**Sample 9 SOH-II-59-01 23-Oct-2024 17:32:32 File: mohammed210_159-9-20241023-1641_COD**

1: MS ES+ :TIC Smooth (SG, 2x3) 2.7e+007

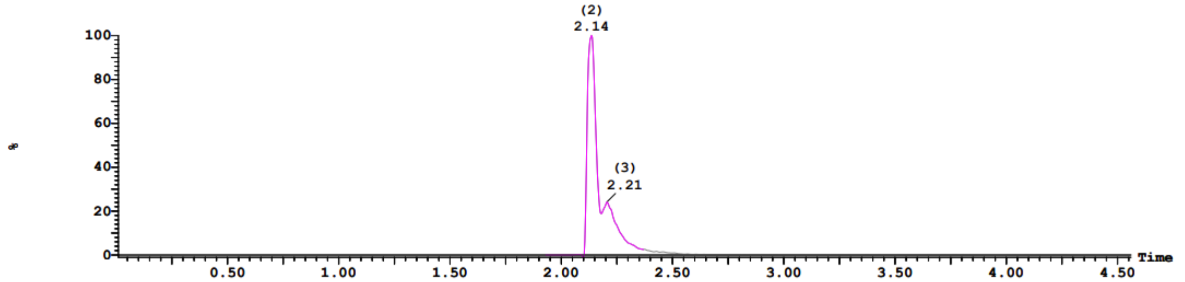


2: UV Detector: TAC: Wavelength Range: (210 - 498) 4.311e+1
Range: 4.311e+1

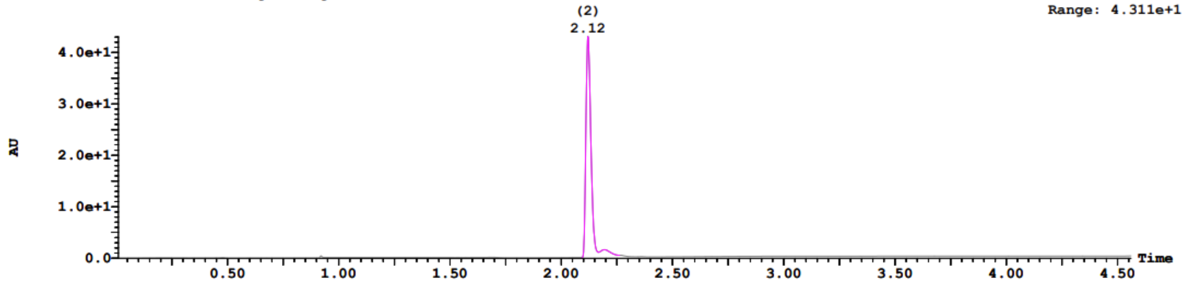


**Sample Report (continued):**

| Peak ID | Time |
|---|---|
| 1 | 2.09 |

1:(Time: 2.09) Combine (297:357-(267:296+358:387)) 1:MS ES+ 3.2e+006

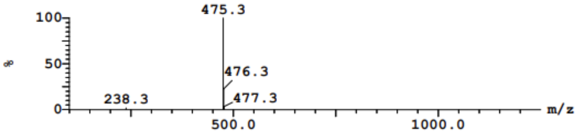


| Peak ID | Time |
|---|---|
| 2 | 2.14 |

2:(Time: 2.12) Combine (306:366-(276:305+367:396)) 1:MS ES+ 4.4e+006

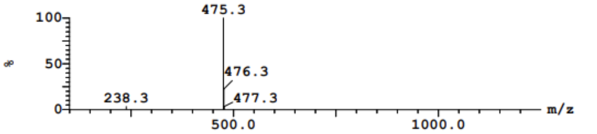


| Peak ID | Time |
|---|---|
| 2 | 2.14 |

2:(Time: 2.12) Combine (1267) 2:UV Detector 9.576e-1 AU

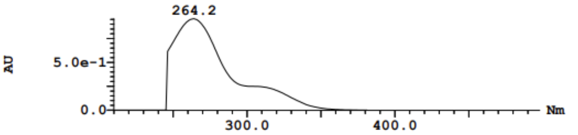


| Peak ID | Time |
|---|---|
| 3 | 2.21 |

3:(Time: 2.21) Combine (332:392-(302:331+393:422)) 1:MS ES+ 6.0e+006

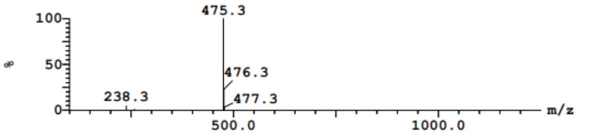


| Peak ID | Time |
|---|---|
| 3 | 2.21 |

3:(Time: 2.19) Combine (1311) 2:UV Detector 5.287e-2 AU

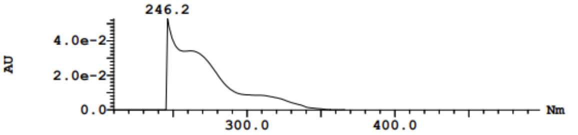


| Peak ID | Time |
|---|---|
| 4 | 2.22 |

4:(Time: 2.22) Combine (336:396-(306:335+397:426)) 1:MS ES+ 5.3e+006

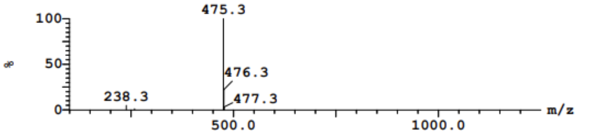
